# Muscle transcriptome profiling reveals novel molecular pathways and biomarkers in laminin-α2 deficient patients

**DOI:** 10.1101/2025.08.23.671372

**Authors:** Veronica Pini, Francesco Catapano, Rosa Bonaccorso, Ben Weisburd, Stefano C. Previtali, Francesco Muntoni

## Abstract

Merosin-deficient congenital muscular dystrophy (LAMA2-RD) is caused by *LAMA2* gene mutations, coding for laminin-211 (merosin) α2 subunit. *LAMA2* mutations leading to complete laminin-211 absence result in an invariably severe clinical phenotype, with profound muscle weakness and respiratory insufficiency. Milder phenotypes are often associated with mutations allowing the production of a partially functional protein. While several dysregulated genes/pathways linked to LAMA2-RD muscle loss are known, an in-depth characterization of LAMA2-RD muscle gene expression profile in patients with mutations differentially affecting *LAMA2* expression is lacking. We generated muscle transcriptomic data from patients with either complete or partial laminin-211 deficiency, and identified pathways linked to the most dysregulated processes. Genes related to fibrosis, inflammation and metabolism were similarly expressed in both patient cohorts. However, a subset of novel pro-fibrotic and pro-inflammatory genes were exclusively expressed in patients (and mice) completely lacking laminin-211, indicating aspects exacerbated in this cohort. Our work characterizes the main contributors of human LAMA2-RD pathology, providing insight into molecular pathways that could be used as disease biomarkers or as targets for therapeutic approaches.

## Introduction

Among the congenital muscular dystrophies (CMDs), merosin-deficient congenital muscular dystrophy (also known as LAMA2-related muscular dystrophy or LAMA2-RD) is considered one of the most severe and common CMD subtypes. It has an estimated global birth prevalence of 8.3 per million live births (Lake *et al*.,2023) and accounts for 30-40% of CMD cases in Europe and the UK (Sframeli *et al*., 2017). LAMA2-RD results from mutations originating in the *LAMA2* gene which codes for the α chain of the heterotrimeric protein laminin-211 (also known as merosin), the main laminin isoform prevalently expressed in the basement membrane of mature skeletal muscles, skin fibroblasts and Schwann cells (Leivo & Engvall, 1988; Sewry *et al*., 1997). Laminin-211 exerts essential structural and signaling roles in maintaining tissue integrity and force transmission by anchoring extracellular matrix (ECM) components to the basal membrane via specific surface receptors, mainly α-dystroglycan and integrin α7β1 (in muscle and fibroblasts) and α6β1 (in Schwann cell). When laminin-211 is absent or defective, these interactions are disrupted, compromising the inside-out and outside-in signaling that normally occurs between ECM and the muscle cell (Mehuron *et al*., 2014). In muscle, the lost or perturbed connection with the extracellular environment disturbs cellular homeostasis, triggering a cascade of molecular events that promote atrophic signaling and ECM rearrangements, lead to muscle fibers degeneration and death, and culminate in their replacement with fibro-adipose tissue (Boppart *et al*., 2011; Han *et al*., 2009).

In a previous genotype-phenotype study we showed that, despite variability, LAMA2-RD severity generally correlates with the extent of laminin-211 expression in muscle (Geranmayeh *et al*., 2010); Patients in whom laminin-211 is completely absent due to loss-of-function mutations in *LAMA2* gene exhibit a severe and clinically uniform congenital phenotype marked by profound muscle weakness, with most affected individuals never achieving independent ambulation (although rare exceptions to this rule have been reported)(Prandini *et al*., 2004). With time, these patients often develop progressive scoliosis and respiratory insufficiency, with most requiring ventilatory support already in the first decade of life. This severe disease course frequently results in premature death, typically by early adulthood, although improvement of standards of care and in particular ventilatory surveillance and intervention continue to make a positive impact on survival. In contrast, hypomorphic *LAMA2* mutations that permit residual protein expression and function lead to a broader and often milder disease spectrum. In this context, muscle weakness typically emerges later and primarily affects limb-girdle muscles, resulting in a phenotype resembling limb-girdle muscular dystrophy (LGMD). Patients expressing partially functional laminin-211 are more likely to maintain ambulation, are less prone to scoliosis, and less frequently require ventilatory support, resulting in longer survival. Myelination defects leading to peripheral neuropathy are also described in both patient cohorts, although the contribution of nerve involvement in human disease appears to be very limited, in contrast to the major role underlying muscle weakness in all murine LAMA2-RD animal models (Previtali & Zambon, 2020).

Differences in clinical severity ascribable to different genotypes and protein levels are not unique to LAMA2-RD; similar genotype-dependent variability was in fact reported also in other neuromuscular conditions such as the dystrophinopathies (where the allelic Duchenne and Becker variants are associated with complete and partial dystrophin deficiency) (J. Wang *et al*., 2021) and the collagenopathies Bethlem-and Ullrich CMD (complete and partial collagen-6 deficiency) (Butterfield *et al*., 2017; Kwong *et al.,* 2023), which share with LAMA2-RD the alteration of molecular aspects such as inflammation and fibrosis, among others.

Although a previous microarray study (Millino *et al*., 2006) highlighted differential expression of select genes in LAMA2-RD patient samples with either complete or partial laminin-α2 deficiency, a comprehensive analysis of all dysregulated molecular pathways in the disease and the global transcriptional landscape of LAMA2-RD muscle across patients with different levels of laminin-α2 expression is currently lacking.

To address this, we analyzed muscle biopsies from unrelated LAMA2-RD patients with either complete or partial laminin-α2 deficiency alongside healthy controls, and performed bulk RNA-sequencing to identify the entire set of genes and molecular traits uniquely dysregulated in these patients-derived samples compared to healthy muscles. Through *in silico* bioinformatic approaches, we defined transcriptomic signatures specific to each patient cohort and identified novel disease biomarkers (some of which are also conserved in LAMA2-RD murine models). This study provides the first exhaustive characterization of the molecular events disrupted by laminin-α2 defects in human muscle, offering new insights into pathogenic mechanisms and identifying novel targets for future therapeutic interventions.

## Results

### RNA profiling of LAMA2-RD muscle samples revealed severity-dependent gene dysregulation in distinct patient cohorts

We extracted total RNA from 12 muscle biopsies, including samples from 4 patients with complete absence of laminin-211 (from now on referred to as the Complete group), 4 patients with partial laminin-211 deficiency (Partial group) and 4 healthy controls, and sequenced them as indicated in the corresponding methods section. Alignment using the STAR algorithm confirmed successful sequencing across all samples, with a mean read alignment rate of 91.75%. ± 2.11 (*Figure 1a*). To assess sample quality and tissue identity, we compared the gene expression profiles of our sequenced samples with muscle- and adipose-specific gene signatures from the Genotype-Tissue Expression (GTEx) project. Three samples (one for each cohort) clustered outside the muscle genes group (*Figure 1b*) and were therefore considered outliers; these were excluded from further bioinformatic analyses. Differentially expressed genes (DEGs) were identified by comparing each patient group to healthy controls using the DESeq2 pipeline. We considered only genes with an adjusted p-value (q-value) ≤ 0.05. The Complete group showed extensive transcriptional dysregulation, with 4,437 DEGs compared to controls (2,517 upregulated and 1,920 downregulated), while the Partial cohort exhibited an intermediate level of dysregulation, with 1,145 DEGs (799 upregulated and 346 downregulated) (*Supplementary Table 1*). These findings support a correlation between the degree of laminin-211 deficiency and the severity of the transcriptional alterations observed in patient-derived muscle tissue.

**Figure 1:**
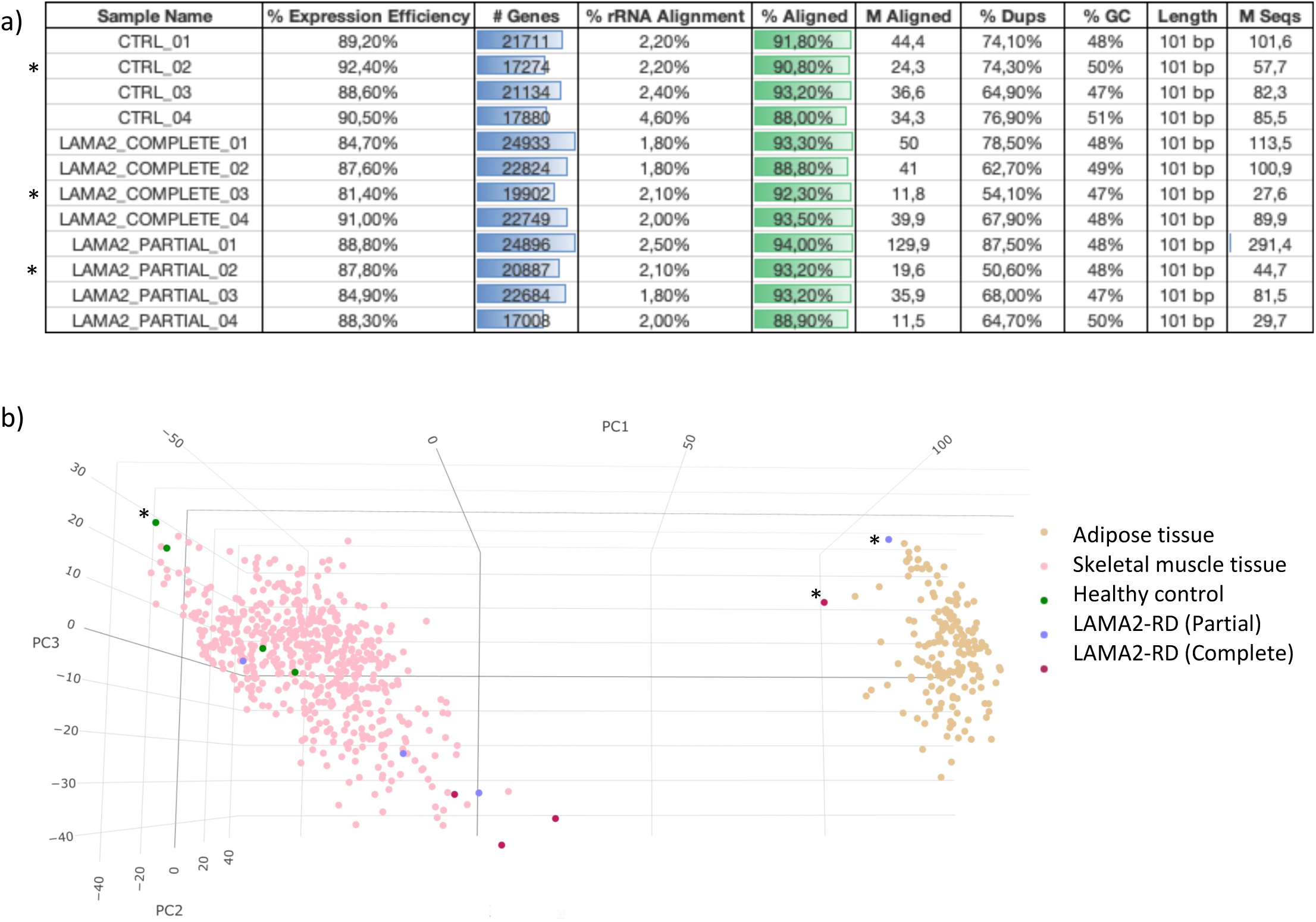
Quality controls run on sequenced samples. a) table including parameters indicative of RNA-sequencing efficiency, including the number of mapped genes, their alignment percentage, the millions (M) of reads sequenced and aligned and the percentage of duplicated regions and GC content. b) principal component analysis plot showing the clustering of sequenced muscle together with genes expressed in skeletal muscle or adipose tissue according to Genotype-Tissue expression (GTEX). * indicates outlier samples.

### DEGs functional enrichment highlighted cohort-specific biological alterations

To identify how dysregulated genes in each patient cohort distribute within the cellular component, molecular function and biological process ontologies (GOs), we performed functional enrichment analysis by running DEGs on the online tool ShinyGO. In the Partial cohort, annotations related to basement membrane, collagen processing and ECM components ranked among the top 20 most significantly enriched terms across all three GO categories, indicating that most dysregulated genes in this group play part in the extracellular remodelling processes associated with the early stages of fibrosis. Similar GO terms were also enriched in the Complete cohort; however, the most significantly enriched annotations in this group referred instead to mitochondrial function and energetic metabolism (*Figure 2, top and middle panels*). To gain a broader overview of the molecular differences between cohorts, we further analysed DEGs against the 50 gene set hallmarks of the Molecular Signatures Database (MSigDB) (*Figure 2, bottom panel*). This analysis revealed that “Oxidative phosphorylation” was the most enriched hallmark in Complete patients (125 genes, Fold Enrichment=3.75), while on the contrary the least enriched set in Partial cohort (22 genes, Fold Enrichment=2,41), indicating the different metabolic compromise between the two. Conversely, “Epithelial Mesenchymal transition (EMT)” hallmark ranked first in Partial (84 genes, Fold Enrichment=9,19) and second in Complete group (121 genes, Fold Enrichment=3.62), suggesting that molecular mechanisms driving the transition towards myofibroblasts are active in both cohorts and are not mitigated by residual laminin-211 expression. Further hallmarks significantly enriched in both cohorts include “inflammatory response”, “myogenesis” “apoptosis”, “hypoxia”, “fatty acids metabolism”, “glycolysis” and signalling pathways associated with T-cells inflammatory response (such as IL2 STAT5 signalling)(Mahmud *et al*., 2013). The full list of significative MSigDB hallmarks in both cohorts, including their fold enrichment and false discovery rate is provided in *Table 1*.

**Figure 2:**
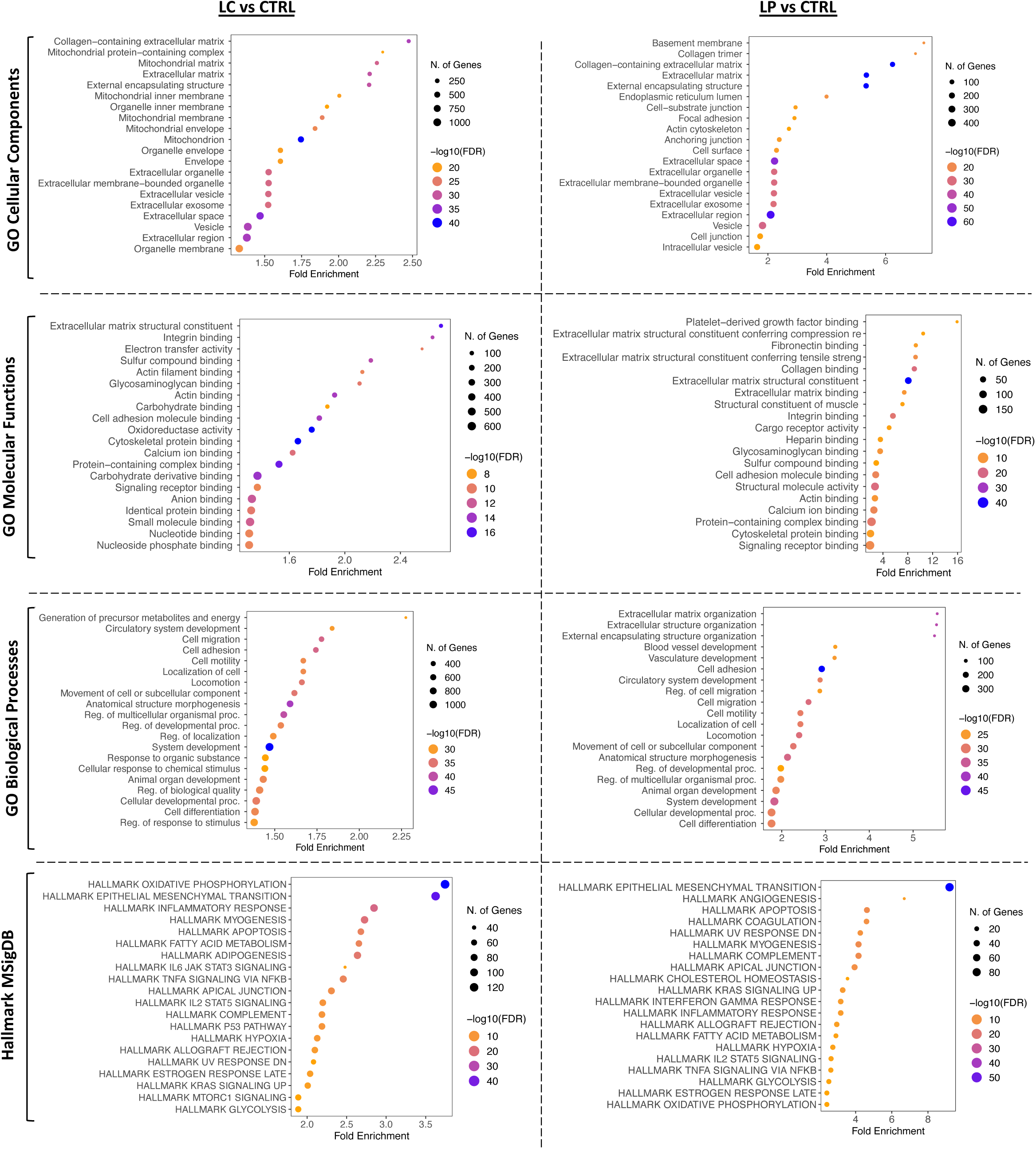
Gene Ontologies and Hallmark gene set Enrichment Analysis of DEGs in separate LAMA2-RD patient cohorts. Dot plots display the 20 most enriched ontologies (GO) populated by genes altered in LAMA2-RD patients with either Complete (LC) or Partial (LP) laminin-211 absence compared to healthy individuals (CTRL). Studied GO categories include cellular components (top panels), molecular functions and biological processes (middle panels). Bottom panels show enrichment of hallmark gene sets from the curated Molecular Signature Databse (MSigDB). Dots size reflects the number of genes associated with each GO term/hallmark, while dots colour varies according to significance, expressed as –log10(FDR). FDR=False Discovery Rate.

**Table 1:** Molecular Signature Database (MSigDB) Hallmarks enriched in skeletal muscle samples of distinct LAMA2-RD patient cohorts. FDR= False Discovery Rate; LC= LAMA2-RD patients with Complete protein deficiency; LP= LAMA2-RD patients with Partial deficiency.

| MSigDB Pathway | Pathway Genes | nGenes LC | Fold Enrichment LC | Enrichment FDR LC | nGenes LP | Fold Enrichment LP | Enrichment FDR LP |
| --- | --- | --- | --- | --- | --- | --- | --- |
| HALLMARK OXIDATIVE PHOSPHORYLATION | 200 | 125 | 3.75 | 1.79E-46 | 22 | 2.41 | 3.15E-04 |
| HALLMARK EPITHELIAL MESENCHYMAL TRANSIT | 200 | 121 | 3.63 | 4.07E-43 | 84 | 9.2 | 2.37E-57 |
| HALLMARK INFLAMMATORY RESPONSE | 200 | 95 | 2.85 | 5.21E-23 | 29 | 3.17 | 1.8E-07 |
| HALLMARK MYOGENESIS | 200 | 91 | 2.73 | 1.53E-20 | 38 | 4.16 | 7.41E-13 |
| HALLMARK ADIPOGENESIS | 200 | 88 | 2.64 | 8.75E-19 | 19 | 2.08 | 4.5E-03 |
| HALLMARK APOPTOSIS | 161 | 72 | 2.68 | 5.39E-16 | 34 | 4.62 | 7.41E-13 |
| HALLMARK TNFA SIGNALING VIA NFKB | 200 | 82 | 2.46 | 1.83E-15 | 24 | 2.63 | 4.77E-05 |
| HALLMARK FATTY ACID METABOLISM | 158 | 70 | 2.66 | 1.99E-15 | 21 | 2.91 | 3.84E-05 |
| HALLMARK APICAL JUNCTION | 77 | 200 | 2.31 | 6.28E-13 | n.a. | n.a. | n.a. |
| HALLMARK IL2 STAT5 SIGNALING | 199 | 73 | 2.2 | 3.82E-11 | 24 | 2.64 | 4.68E-05 |
| HALLMARK P53 PATHWAY | 200 | 73 | 2.19 | 4.2E-11 | 18 | 1.97 | 9.47E-03 |
| HALLMARK COMPLEMENT | 200 | 73 | 2.19 | 4.2E-11 | 38 | 4.16 | 7.41E-13 |
| HALLMARK HYPOXIA | 200 | 71 | 2.13 | 3.23E-10 | 25 | 2.74 | 1.91E-05 |
| HALLMARK ALLOGRAFT REJECTION | 200 | 70 | 2.1 | 8.37E-10 | 27 | 2.96 | 1.92E-06 |
| HALLMARK ESTROGEN RESPONSE LATE | 200 | 68 | 2.04 | 5.7E-09 | 22 | 2.41 | 3.15E-04 |
| HALLMARK KRAS SIGNALING UP | 200 | 67 | 2.01 | 1.39E-08 | 30 | 3.28 | 5.95E-08 |
| HALLMARK IL6 JAK STAT3 SIGNALING | 87 | 36 | 2.48 | 1.26E-07 | 10 | 2.52 | 1.14E-02 |
| HALLMARK UV RESPONSE DN | 144 | 50 | 2.08 | 3.05E-07 | 28 | 4.26 | 4.82E-10 |
| HALLMARK MTORC1 SIGNALING | 200 | 63 | 1.89 | 4.19E-07 | n.a. | n.a. | n.a. |
| HALLMARK GLYCOLYSIS | 200 | 63 | 1.89 | 4.19E-07 | 23 | 2.52 | 1.32E-04 |
| HALLMARK ANGIOGENESIS | 36 | 19 | 3.16 | 1.8E-06 | 11 | 6.69 | 1.6E-06 |
| HALLMARK INTERFERON GAMMA RESPONSE | 200 | 61 | 1.83 | 2.04E-06 | 29 | 3.17 | 1.8E-07 |
| HALLMARK PEROXISOME | 104 | 36 | 2.07 | 1.48E-05 | 10 | 2.11 | 3.35E-02 |
| HALLMARK APICAL SURFACE | 44 | 20 | 2.72 | 1.62E-05 | n.a. | n.a. | n.a. |
| HALLMARK COAGULATION | 138 | 43 | 1.87 | 3.86E-05 | 29 | 4.6 | 3.43E-11 |
| HALLMARK CHOLESTEROL HOMEOSTASIS | 74 | 27 | 2.19 | 6.16E-05 | 12 | 3.55 | 3.15E-04 |
| HALLMARK XENOBIOTIC METABOLISM | 200 | 55 | 1.65 | 1.4E-04 | n.a. | n.a. | n.a. |
| HALLMARK ESTROGEN RESPONSE EARLY | 200 | 55 | 1.65 | 1.4E-04 | 21 | 2.3 | 7.65E-04 |
| HALLMARK BILE ACID METABOLISM | 112 | 35 | 1.87 | 1.78E-04 | n.a. | n.a. | n.a. |
| HALLMARK UV RESPONSE UP | 158 | 45 | 1.71 | 2.35E-04 | 15 | 2.08 | 1.12E-02 |
| HALLMARK HEME METABOLISM | 200 | 51 | 1.53 | 1.56E-03 | n.a. | n.a. | n.a. |
| HALLMARK HEDGEHOG SIGNALING | 36 | 13 | 2.16 | 6.1E-03 | n.a. | n.a. | n.a. |
| HALLMARK NOTCH SIGNALING | 32 | 11 | 2.06 | 1.71E-02 | n.a. | n.a. | n.a. |
| HALLMARK REACTIVE OXYGEN SPECIES PATHW | 49 | 15 | 1.83 | 1.71E-02 | n.a. | n.a. | n.a. |
| HALLMARK PI3K AKT MTOR SIGNALING | 105 | 27 | 1.54 | 1.74E-02 | 11 | 2.29 | 1.49E-02 |
| HALLMARK ANDROGEN RESPONSE | 100 | 25 | 1.5 | 3.02E-02 | 14 | 3.07 | 4.16E-04 |
| HALLMARK INTERFERON ALPHA RESPONSE | 97 | n.a. | n.a. | n.a. | 10 | 2.26 | 2.22E-02 |
| HALLMARK MITOTIC SPINDLE | 199 | n.a. | n.a. | n.a. | 18 | 1.98 | 9.36E-03 |
| HALLMARK APICAL JUNCTION | 200 | n.a. | n.a. | n.a. | 36 | 3.94 | 1.46E-11 |

### Core pathway and upstream regulator analyses indicated directional regulation of transcriptional cascades within LAMA2-RD muscle

While highlighting key molecular mechanisms affected by laminin-211 dysregulation, enrichment analysis did not provide insights into the directionality (activation or inhibition) of the above-mentioned dysregulated processes. To address this, we conducted a more granular analysis relying on Ingenuity Pathway Analysis (IPA), a knowledge-based software able to infer pathways activation state. For each cohort, we analysed DEGs against findings observed in skeletal muscle-resident cell types (such as myoblasts, fibroblasts, immune cells, adipocytes and endothelial cells) as well as cells from adjacent tissues which could contribute to muscle biopsy heterogeneity (smooth muscle, motor neurons and chondrocytes) (MB filters).

In the complete cohort, altered canonical pathways (CP) were assigned to 40 macro categories that confirmed gene enrichment data and highlighted profound alterations in energy metabolism, oxidative stress, inflammation, and fibrosis. These findings inferred a downregulation of energy metabolism (reflected by decreased expression of the transcription factors *PPARGC1A, PPARA, PPARG*, and *KLF15*) and an upregulation of oxidative stress, inflammation, and fibrosis processes (mainly increased expression of *TNF, IL6, IL1B*, and *IFNG*) (*Figure 3a* and *Supplementary Table 2*). When ranked for -log(p-value), among CPs for which activation status was predicted the most significantly affected CPs were related to cellular metabolism and mitochondrial dysfunction, especially ATP production and oxidative phosphorylation (both markedly downregulated, z-=-8.19 and z=-7.35) (*Figure 3b*), followed by overactive CPs connected to multiple inflammatory and hypoxic processes (involving sirtuins, neutrophils, lymphoid cells, cytokines and activated platelets signalling) as well as tissue fibrosis and extracellular matrix remodelling (collagens, elastic fibres). Importantly, integrin and muscle contraction pathways, closely associated with laminin-211’s role in muscle structure and mechanotransduction, were also among the most significantly dysregulated. Overlapping Canonical Pathways data analysis revealed that, generally, upregulated genes contributed to exacerbate more than one signalling pathway transversely supporting both fibrosis and inflammation, while a distinct pool of (mostly) downregulated genes contributed uniquely to perturb energetic homeostasis (*Figure 3c*). When analysing upstream regulators, priority was given to transcription factors (TFs) and miRNAs, given their potential as therapeutic targets. Among the 5 most downregulated TFs *PPARGC1A* was not only the most significant (p-value=2.24E-35), but also the one with the lowest z-score (z=-8.42), followed by the cell-cycle regulators *RB1* (z=-4.84), *GFI1* (z=-4.39) and *SMARCA5* (z=-4.25) and finally *KLF15* (z=-3.58) which, besides supporting many metabolic pathways, also contributes to myoblasts differentiation and regeneration (Gao *et al*., 2023). The highest z-score was instead assigned to *NFKBIA* (z=+5.74), *TP53* (z=+5.73) and *BHLHE40* (z=+5.50), involved in cytokine release and cell death, together with *KLF6* (z=+5.59) and *SMAD3* (z=+5.45) that are generally overexpressed in fibrotic milieu. Upregulated miRNAs identified as upstream regulators included mir-223 (z=+3.39), which promotes granulocyte and megakaryocyte generation influencing myoblasts proliferation and differentiation (Li *et al*., 2017) (Cheng *et al*., 2020) and miR-21 (z=+1.91), which upregulation supports atrophy and fibrosis acting on the TGF-β smad2/3 pathway (X. Song *et al*., 2024). The anti-fibrotic mir-146 (Sun *et al*., 2017) was instead assigned a negative z-score (z=-1.71), together with mir-140 (z=-1.99) and mir-155 (z=-2.07), both acting on regeneration and inflammation and whose role in skeletal muscle is still controversial (Lopes *et al.,* 2024; Nie *et al.,* 2016; Pan *et al.,* 2023) (*Figure 3d*). The complete list of dysregulated upstream regulators is provided in *Table 2*.

**Figure 3:**
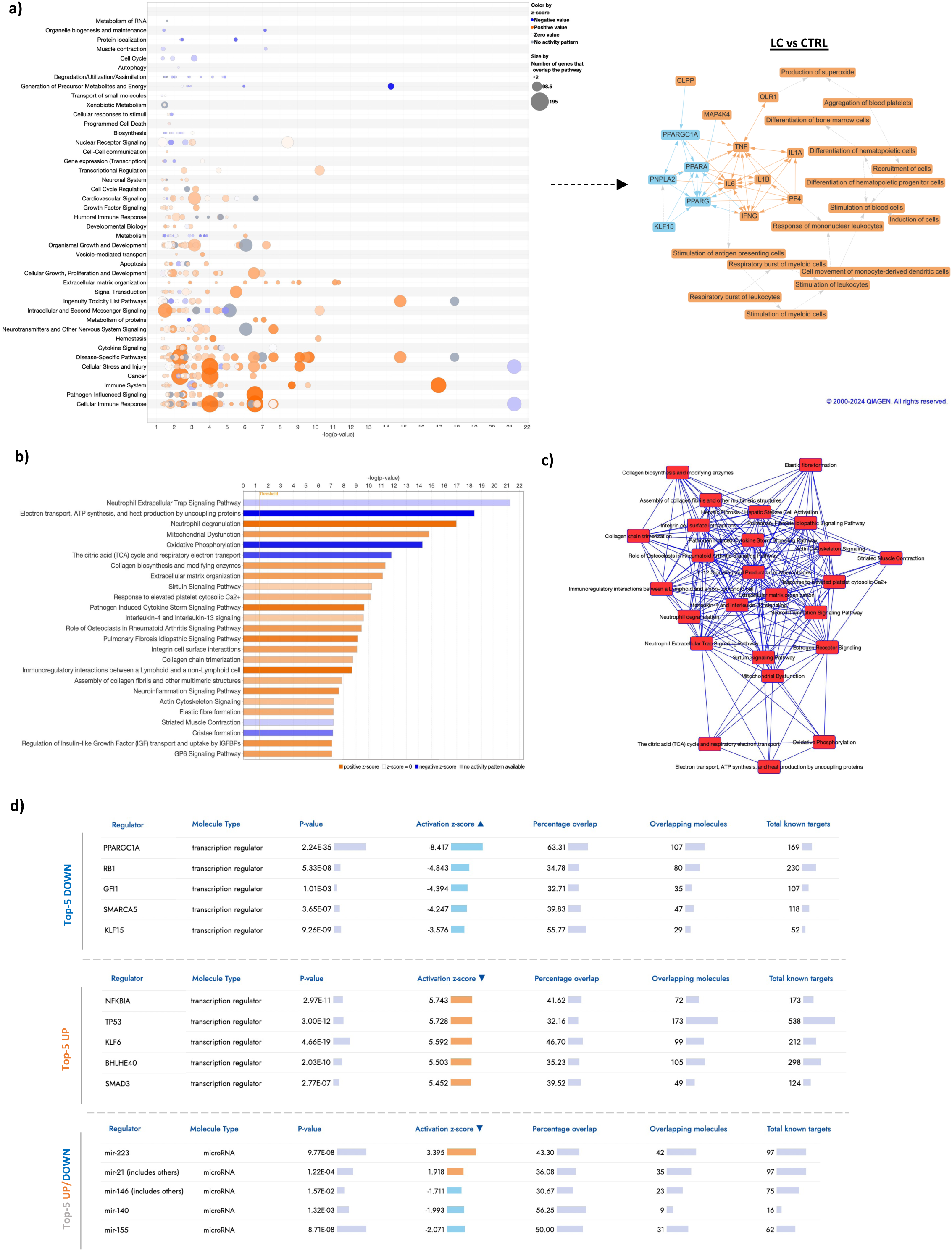
IPA analysis of Canonical Pathways and Regulators altered in Complete patient cohort (LC) a) bubble chart showing macro-categories of altered pathways and graphical summary of DEGs connection; b) top 25 dysregulated pathways ranked by statistical significance; c) pathways sharing one or more genes; d) upstream regulators including the ten most perturbed transcription factors and altered miRNAs.

**Table 2:** IPA upstream regulators in Complete cohort. Transcription factors and miRNA listed in the table are ordered according to p-value of overlap with DEGs in this cohort.

| Upstream Regulator | Molecule Type | Predicted Activation State | Activation z-score | p-value of overlap |
| --- | --- | --- | --- | --- |
| PPARGC1A | transcription regulator | Inhibited | -8.417 | 2.24E-35 |
| KLF6 | transcription regulator | Activated | 5.592 | 4.66E-19 |
| MAFB | transcription regulator | Activated | 3.219 | 9.1E-13 |
| ARID1A | transcription regulator | Activated | 3.37 | 1.65E-12 |
| TP53 | transcription regulator | Activated | 5.728 | 3E-12 |
| NFKBIA | transcription regulator | Activated | 5.743 | 2.97E-11 |
| STAT3 | transcription regulator | Activated | 3.451 | 7.04E-11 |
| BHLHE40 | transcription regulator | Activated | 5.503 | 2.03E-10 |
| YAP1 | transcription regulator | Activated | 3.334 | 2.97E-09 |
| NRIP1 | transcription regulator | Activated | 4.36 | 5.87E-09 |
| KLF15 | transcription regulator | Inhibited | -3.576 | 9.26E-09 |
| H2AX | transcription regulator | Inhibited | -2.777 | 4.86E-08 |
| RB1 | transcription regulator | Inhibited | -4.843 | 5.33E-08 |
| SMAD3 | transcription regulator | Activated | 5.452 | 0.000000277 |
| ID3 | transcription regulator | Activated | 2.106 | 0.00000033 |
| SMARCA5 | transcription regulator | Inhibited | -4.247 | 0.000000365 |
| SOX4 | transcription regulator | Activated | 3.326 | 0.000000874 |
| BCL6 | transcription regulator | Inhibited | -3.993 | 0.00000127 |
| IFI16 | transcription regulator | Activated | 2.931 | 0.00000134 |
| HMGB1 | transcription regulator | Activated | 3.883 | 0.00000238 |
| FOXO3 | transcription regulator | Inhibited | -2.345 | 0.00000352 |
| BRCA1 | transcription regulator | Activated | 2.551 | 0.00000838 |
| SMAD7 | transcription regulator | Inhibited | -3.04 | 0.00000892 |
| CTNNB1 | transcription regulator | Activated | 2.297 | 0.0000136 |
| ZBTB7B | transcription regulator | Inhibited | -2.469 | 0.0000255 |
| TBX5 | transcription regulator | Inhibited | -2.321 | 0.0000448 |
| FOXO4 | transcription regulator | Inhibited | -2.15 | 0.000054 |
| IKZF1 | transcription regulator | Inhibited | -3.52 | 0.0000543 |
| NFKB1 | transcription regulator | Activated | 2.46 | 0.0000853 |
| MYOCD | transcription regulator | Inhibited | -2.047 | 0.0000904 |
| KLF11 | transcription regulator | Inhibited | -2.53 | 0.0000988 |
| VDR | transcription regulator | Activated | 3.772 | 0.000107 |
| NFIC | transcription regulator | Activated | 2.324 | 0.000118 |
| PPARGC1B | transcription regulator | Inhibited | -2.087 | 0.000192 |
| HDAC2 | transcription regulator | Inhibited | -2.236 | 0.000201 |
| TAF4 | transcription regulator | Inhibited | -2.606 | 0.000442 |
| KLF2 | transcription regulator | Inhibited | -3.269 | 0.000451 |
| MRTFA | transcription regulator | Activated | 4.123 | 0.000485 |
| ATF3 | transcription regulator | Inhibited | -3.004 | 0.000579 |
| GPS2 | transcription regulator | Inhibited | -2.354 | 0.000758 |
| STAT1 | transcription regulator | Activated | 3.363 | 0.000825 |
| MRTFB | transcription regulator | Activated | 3.491 | 0.000929 |
| ETS1 | transcription regulator | Activated | 3.018 | 0.000973 |
| GFI1 | transcription regulator | Inhibited | -4.394 | 0.00101 |
| MEOX2 | transcription regulator | Inhibited | -2.683 | 0.00149 |
| EGR1 | transcription regulator | Activated | 3.242 | 0.00209 |
| TCF3 | transcription regulator | Activated | 2.483 | 0.00269 |
| NRF1 | transcription regulator | Inhibited | -2.646 | 0.00288 |
| MEF2D | transcription regulator | Activated | 2.376 | 0.00363 |
| GATA4 | transcription regulator | Inhibited | -2.615 | 0.00401 |
| RELA | transcription regulator | Activated | 3.512 | 0.00408 |
| ZFP36 | transcription regulator | Inhibited | -2.483 | 0.00443 |
| EZH2 | transcription regulator | Inhibited | -2.022 | 0.00527 |
| SPIB | transcription regulator | Activated | 3.714 | 0.00545 |
| TSC22D1 | transcription regulator | Activated | 2 | 0.00635 |
| GLI1 | transcription regulator | Activated | 3.543 | 0.00746 |
| IRF8 | transcription regulator | Activated | 2.494 | 0.0084 |
| BTG2 | transcription regulator | Inhibited | -2.961 | 0.00903 |
| PML | transcription regulator | Activated | 2.386 | 0.0113 |
| TAF4B | transcription regulator | Activated | 2 | 0.0161 |
| KLF5 | transcription regulator | Activated | 2.433 | 0.0181 |
| FOXC1 | transcription regulator | Activated | 2.309 | 0.0193 |
| GLI3 | transcription regulator | Activated | 2.111 | 0.0193 |
| E2F1 | transcription regulator | Activated | 3.07 | 0.0257 |
| NFATC1 | transcription regulator | Activated | 2.397 | 0.0264 |
| NFKBIE | transcription regulator | Activated | 2 | 0.0309 |
| VENTX | transcription regulator | Activated | 2 | 0.0317 |
| IKZF4 | transcription regulator | Activated | 2.813 | 0.0398 |
| RUNX2 | transcription regulator | Activated | 2.209 | 0.045 |
| EPAS1 | transcription regulator | Activated | 2.75 | 0.0476 |
| mir-223 | microRNA | Activated | 3.395 | 9.77E-08 |
| mir-21 (includes other) | microRNA | Activated | 1.918 | 0.000122 |
| mir-146 (includes other) | microRNA | Inhibited | -1.711 | 0.0157 |
| mir-140 | microRNA | Inhibited | -1.993 | 0.00132 |
| mir-155 | microRNA | Inhibited | -2.071 | 8.71E-08 |

© 2000-2025 QIAGEN. All rights reserved.

A partially different scenario emerged from the Core analysis run in Partial cohort, for which IPA predicted the exacerbation of pathways mainly associated with *TGFB1* expression; within the 35 affected macro categories, metabolic pathways were in fact less enriched, whereas pathways related to ECM remodelling, inflammatory cytokine production, and immune responses were similarly represented as in the Complete group (*Figure 4a*). The analysis of the top 25 most significant pathways confirmed the different perturbation trend in the two cohorts and the negligible impact of partial protein deficiency on mitochondrial-metabolic genes (*Figure 4b and 4c*), suggesting that even reduced levels of laminin-211 may partially preserve metabolic integrity. Upstream regulator analysis *(Figure 4d*) anticipated *PPARGC1A* to be the most significantly downregulated TF in this cohort too (z-score=-6.03), followed by *IKZF1* (z-score=-3.28), *KLF2* (z-score=-3.00), *ETV3* (z-score=-3.00), and *TBX5* (z-score=-2.99), mainly implicated in controlling T and B-cells development and pro-fibrotic inflammation (Chrysanthopoulou *et al*., 2023; Villar *et al*., 2023). As for Complete cohort, the 5 most upregulated transcription factors included *NFKB1A* (z-score=+5.36), *SMAD3* (z-score=+5.22), *TP53* (z-score=+5.20) and *BHLHE40* (z-score=+3.90) with the sole difference being *MRTFA* (z-score=+4.05), actively involved in epithelial-mesenchymal transition (Bialik *et al*., 2019). Finally, miRNAs predicted to be dysregulated included mir-21 (z=+2.26), mir-223 (z=+1.51) and mir-155 (z=-1.66), consistent with findings in Complete cohort. The downregulation of 3 further miRNA was linked to increased EMT transition and Sonic Hedgehog signalling (mir-506, z=-1.99) (Haque *et al*., 2020), increased fibrosis and inflammation (let-7, z=-2.21) (M. Zhu *et al*., 2019) (J. Song *et al*., 2023) and muscle atrophy/wasting (mir-29, z=-3.09) (*Figure 4d and Table 3*).

**Figure 4:**
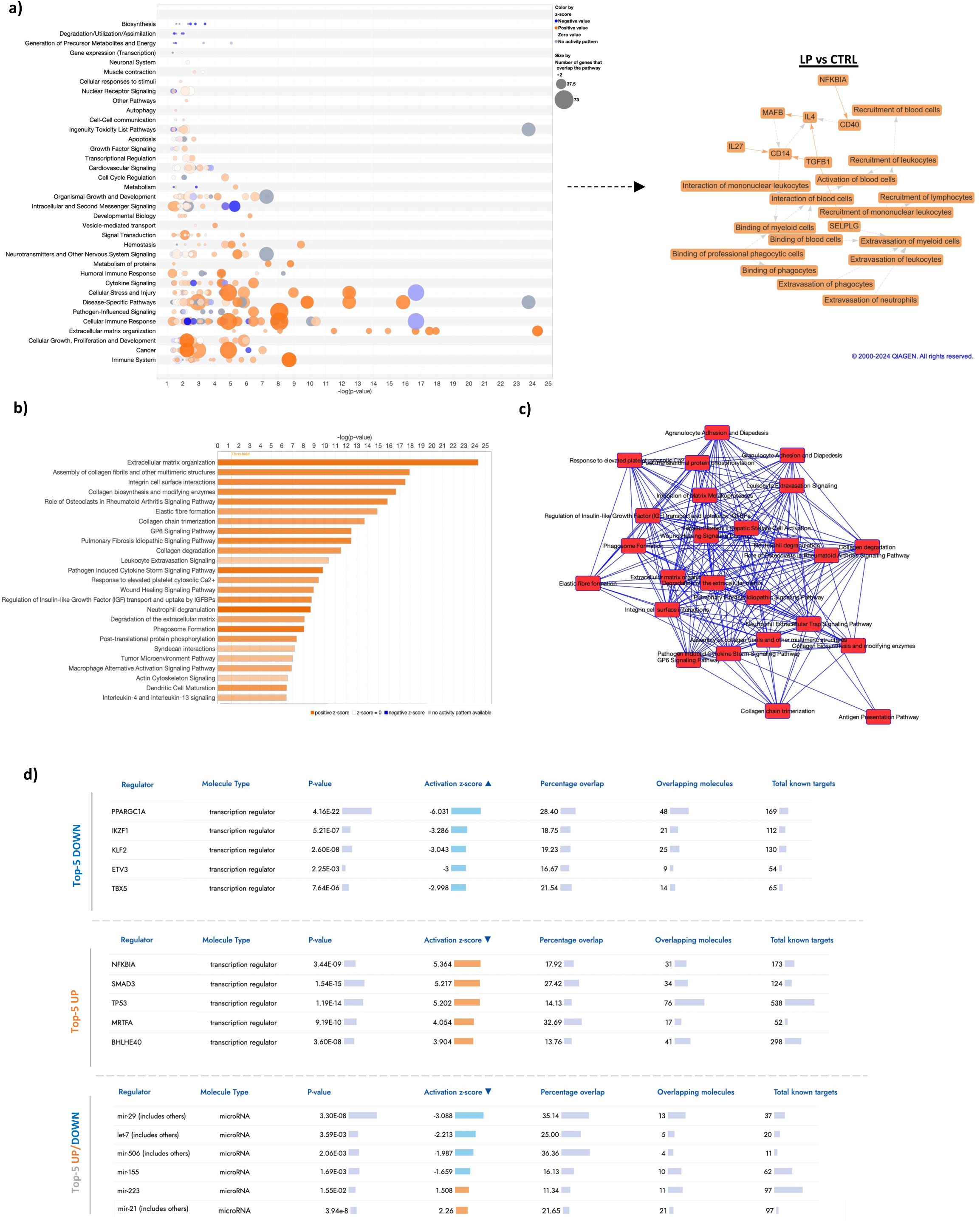
IPA analysis of Canonical Pathways and Regulators altered in Partial patient cohort (LP) a) bubble chart showing affected macro-categories and graphical summary of DEGs connection; b) top 25 altered pathways ranked by statistical significance; c) pathways sharing one or more genes; e) upstream regulators including the ten most perturbed transcription factors and altered miRNAs.

**Table 3:**
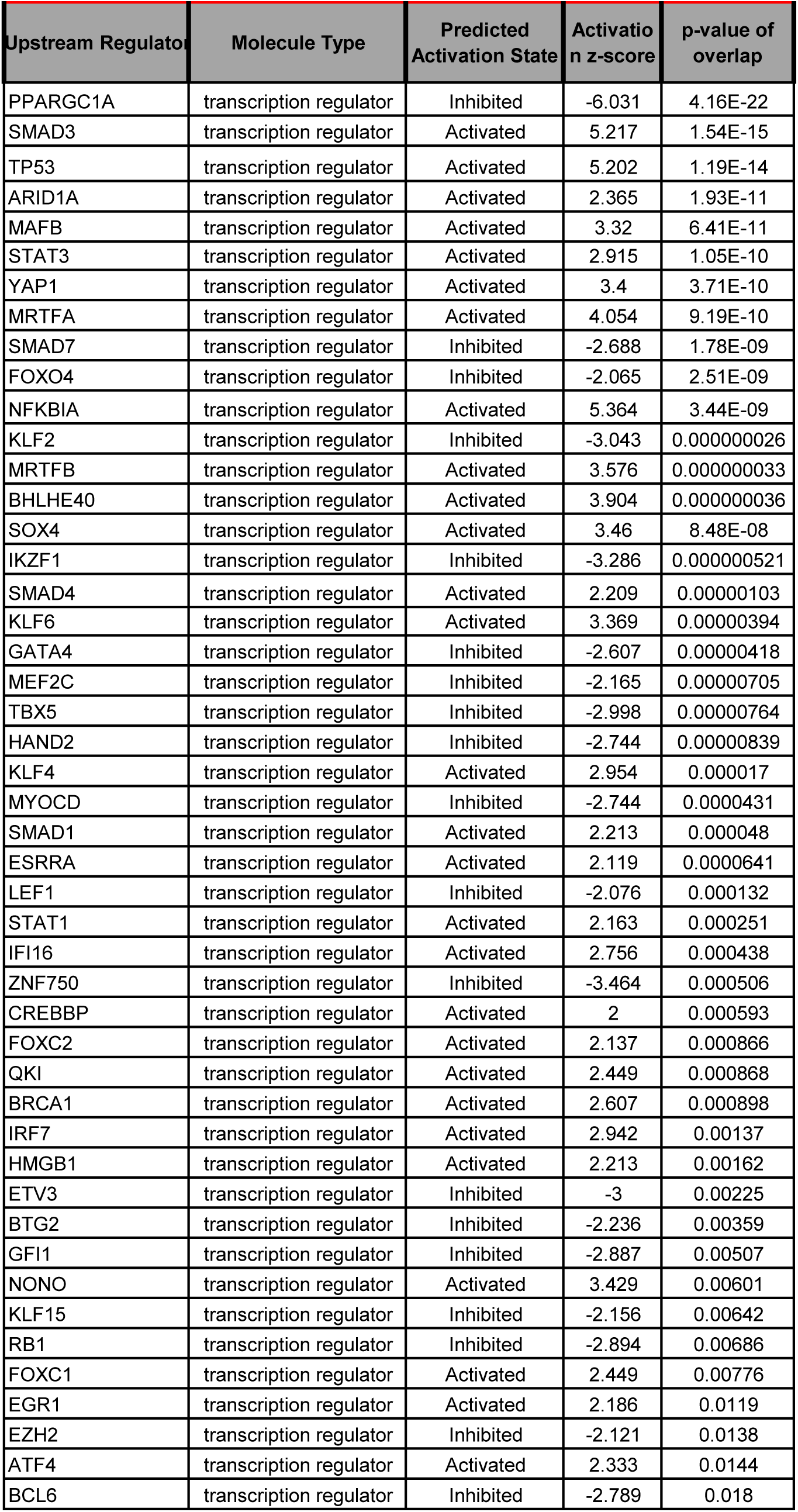
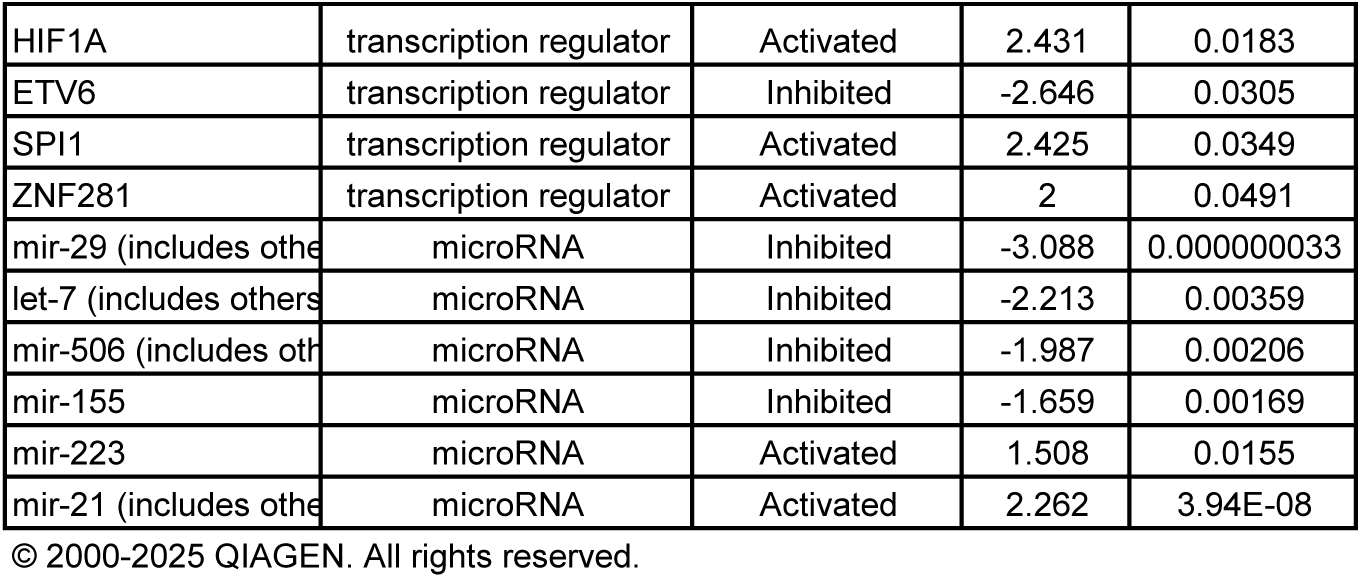
IPA upstream regulators in Partial cohort. Transcription factors and miRNA listed in the table are ordered according to p-value of overlap with DEGs in this cohort.

### Comparative analysis highlighted context-dependent dysregulation in LAMA2-RD muscle and emphasized processes uniquely altered in Complete patient cohort

To gain a deeper understanding of the variability between the two cohorts and identify secondary pathways involved in disease pathogenesis, we conducted a comparative analysis of dysregulated CPs, ranking them for their z-score value (indicative of their perturbation extent) in the more compromised Complete cohort. This approach facilitated the identification of CPs uniquely perturbed in the Complete group, shown in *Supplementary Figure 1*. Our examination was then restricted to the top 100 altered processes with a z-score cut-off of 1.5 in at least one cohort, focusing on the 50 most active and 50 most downregulated CP. Generally, CPs with the highest significance were also the most down/up regulated; except for a limited number of CPs with a nearly identical z-score, most CPs altered in both cohorts showed stronger perturbation in the Complete group (*Figure 5*). Analysing the Top50 downregulated CPs, we gathered further insights into the widespread metabolic dysfunction in LAMA2-RD patients, extended beyond lipids metabolism to include carbohydrates processing and amino-acids catabolism (*Figure 5, left panel*). The highest z-score differences between cohorts included mitochondrial pathways, with “Electron transport and ATP synthesis” and “Oxidative phosphorylation” being the most suppressed in the Complete group. Mitochondrial functions such biogenesis, translation and peroxisomal protein import were also severely compromised in Complete cohort but only slightly impacted (z-score <-2) in patients with partially functional laminin. Remarkably, 22 of the 50 downregulated pathways uniquely affected in patients with complete protein absence were primarily associated to mitochondria morphology and function (such as “Cristae formation” “Carnitine metabolism”, “NAD signalling”), processing of amino acids/cofactors and impaired skeletal muscle contraction. In contrast, z-score differences in the Top50 upregulated CPs were generally less pronounced between cohorts (*Figure 5, left panel*). “Phagosome formation” showed the highest z-score (z=+8.21) in Complete patients, alongside several CPs suggestive of chronic inflammation following tissue injury (Howard *et al*., 2021). FAK signalling was also dramatically upregulated (z-score > +6 in both cohorts) potentially compensating for laminin-211 loss by promoting cell survival, but eventually also contributing to fibroblasts activation and tissue remodelling (Zhao *et al*., 2016). A small subset of CPs with a z-score difference > 2 between cohorts highlighted novel CPs not yet (or poorly) studied in muscular dystrophies that may explain the more severe molecular signature of Complete LAMA2-RD patients. These include pathways active in inflammatory environments (“HOTAIR”(Chini *et al*., 2023) and “TREM1 signalling” (Howard *et al*., 2021)), inflammation-related cell death mechanisms (“Granzyme A signalling” (Zhu *et al.,* 2009) and “Pyroptosis” (Dubuisson *et al.,* 2022)), and CPs implicated in nervous system perturbation (“Synaptogenesis” and “Myelination signalling”)(Shorer *et al*., 1995; Tian *et al*., 1997). Finally, to pinpoint how pathway perturbations varied across cohorts under different filtering conditions, we performed a series of increasingly stringent analyses, sequentially excluding one cell type at a time; we then compared each analysis to the original transcriptional signature obtained with MB constraints. This approach revealed that most pathways predicted to be upregulated in the Complete cohort were less perturbed when analyses were restricted to single muscle-specific cell types, suggesting that the interplay of multiple cell populations within the muscle microenvironment plays a crucial role in exacerbating those mechanisms. In contrast, filtering had minimal impact on downregulated pathways in Complete group, and both on up-and downregulated pathways in Partial cohort too (*Supplementary Figure 2 and Supplementary Table 3*). These findings highlighted the importance of considering cellular context when interpreting transcriptomic data, particularly in heterogeneous tissues such as skeletal muscle.

**Figure 5:**
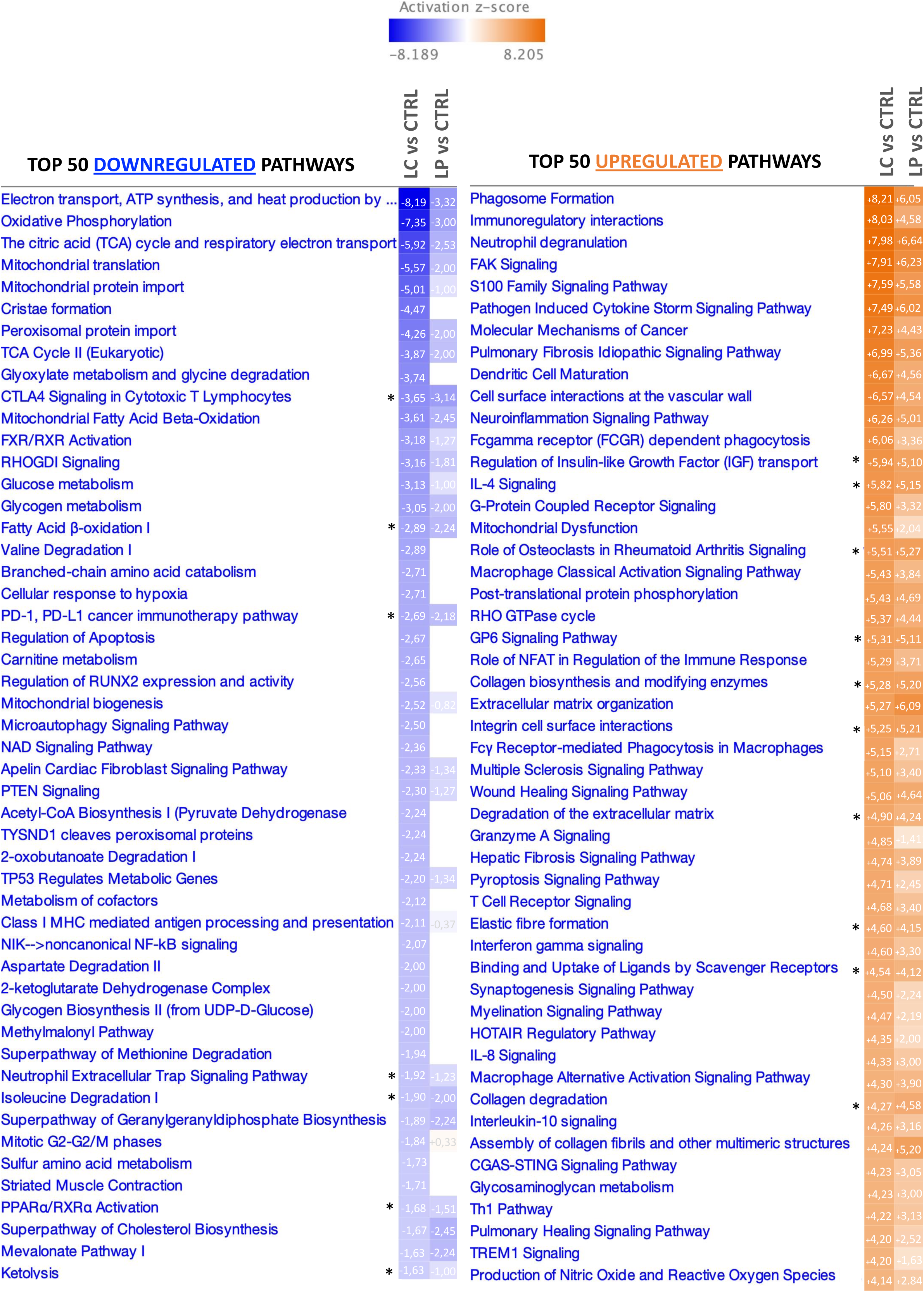
Comparison of top 100 altered pathways identified by IPA in both patient cohorts. Top 50 downregulated pathways (left panel) and Top 50 upregulated pathways (right panel) are ranked according to their activation z-score in Complete patients (LC). LP=Partial cohort. Z-score values for each cohort are indicated aside pathways name. *=similarly altered pathways

### Targeted DEGs analysis Identified core genes linked to key pathogenic mechanisms in LAMA2-RD

In the final stage of our study, we performed a focused gene-by-gene analysis of the transcriptomic data to better define the contribution of individual genes to the most dysregulated biological processes. Supported by literature evidence on DEGs, we could associate 1061 genes altered in Complete cohort with known LAMA2-RD traits; these included fibrosis, ECM remodelling, inflammatory processes, energetic metabolism and oxidative stress response, protein processing/turnover, cell death, cell cycle and muscle contraction, impaired organelles homeostasis in sarcoplasm, and disrupted crosstalk with motor neuron (*Supplementary Table 4*). As expected considering cohort-specific differences found in each of the above-mentioned analyses, only 359 genes were also dysregulated in Partial cohort, and generally to a lesser extent compared to the Complete cohort. The 139 fibrosis/ECM-related genes were similarly altered in the two groups, except for a few genes (including *SPP1, POSTN* and *GPC3)* only altered in Complete. In contrast, the inflammatory response appeared markedly more pronounced in patients completely deficient in laminin-211, as out of 211 inflammation-related genes only 77 were also altered in Partial cohort. Within this category, overexpressed DEGs unique to Complete cohort included those coding for immunoglobulins and inflammatory markers (*PPBP, PF4, TREM2*), T-cells, leukocytes, interleukins, and inflammasome components. Genes related to metabolic imbalance also represented a substantial category (n=304), yet only 69 were dysregulated in both cohorts. Most of the perturbed DEGs in Complete were linked to mitochondrial biogenesis (*PERM1, PPARGC1B*, and *PPTC7*) and mitochondrial protein synthesis/processing (*MTFR1*), together with genes coding for key proteins for lipids metabolism including β-oxidation and electron transport (*NDUFA9, NDUFS2*). A higher number of genes linked to glycogen (*HK2*) and glucose (*PFKB3*) metabolism were also downregulated in the Complete cohort contributing to the energetic deficit, similar to what we found for genes involved in metabolism of amino acids (*GOT1, BCAT1*), nucleotides (*GMPR, NT5C1A*) and vitamins (*PANK1, RETSAT*). Genes linked with cellular stress response followed the same trend (i.e. out of 183 genes, only 57 shared between cohorts, including *PERP, TPRG1* and *SOD3*). We inferred heightened oxidative stress in Complete patients by the strong upregulation of *CDH22* and *THRA4*, *GSTM5, MT1A, NRROS,* among others, and the consequent alteration of key regulators of processes like cell death, autophagy and ubiquitin-proteasome response. Notably, among genes associated with this degradation pathway, most genes were slightly downregulated apart from *UBASH3A* which was strongly upregulated. A small number of mostly downregulated DEGs supporting structural stability and sarcomere organization (n=76, of which 27 altered in both cohorts) likely reflected the disrupted laminin-α-dystroglycan axis. Within these, both patient cohorts exhibited high expression of *MYH8* and *MYH3* gene, suggesting active muscle regeneration. An even smaller number of genes (n=50) linked to multiple cell cycle phases showed differential expression, with only 10 including *CDKN1A, GADD45A* and *RPRM* (the most upregulated) common to both groups. We also identified further 85 genes (of which 26 altered in both Cohort) related to neuromuscular junction/transmission of nervous signal, which we attributed to the presence of motor nerve terminals in the processed muscle biopsy. Interestingly, these were mostly overexpressed, suggesting a potential compensatory response to impaired nerve-muscle communication. Finally, 13 genes were linked to altered signalling pathways such as Notch, Hedgehog and Wnt.

### Validation of selected DEGs in LAMA2-RD patient-and dystrophic mice-derived tissue by qPCR shed light on novel disease biomarkers

Among DEGs we identified a limited number of overexpressed genes coding for protein (*OSTN, COL22A1, COMP, ARGHAP36, PF4, NLRP3*) and lncRNA *(HOTAIR, XIST)* whose function is linked to general pathological aspects occurring in dystrophic muscles (i.e. fibrosis and inflammation); we further validated their expression by qPCR on RNA extracted from sequenced tissues. Except for *NLRP3* (Jeudy *et al*., 2011), none of these genes was previously studied in relation to LAMA2-RD or other muscular dystrophies*. OSTN,* coding for the bone-specific secreted protein osteocrin, was selected as it was one of the most upregulated gene in both cohorts (LC=+10,68 and LP=+11,05), followed by *COMP* (LC=+9,43 and LP=+6,80), *COL22A1* (LC=+8,11, LP=+8,02) and *ARHGAP36* (LC=+8,08, LP=+5,91), involved in the production of ECM components. *PF4*, recently identified as a key molecule concomitantly regulating inflammation and fibrosis (Silva-Cardoso *et al*., 2020) was selected for its prominent overexpression (LC=+7,05) in Complete cohort only, similarly to the less perturbed *NLRP3* (+2,64), a key component of the inflammasome also exacerbated in DMD (Boursereau *et al*., 2018). NLRP3 acts as a downstream effector of the lncRNA *HOTAIR*, also uniquely dysregulated in Complete cohort (LC=2,16). Finally, we considered the epigenetic regulator *XIST* as it was not only one of the most upregulated (LC=+10.09) genes uniquely altered in Complete cohort, but also as the most significant (q-value= 5,83E-65) within DEGs. Expression data of tested genes generally concurred with what detected by RNA-sequencing, although with variable significance and more pronounced fold changes measured by qPCR (*Figure 6, left panel*).

**Figure 6:**
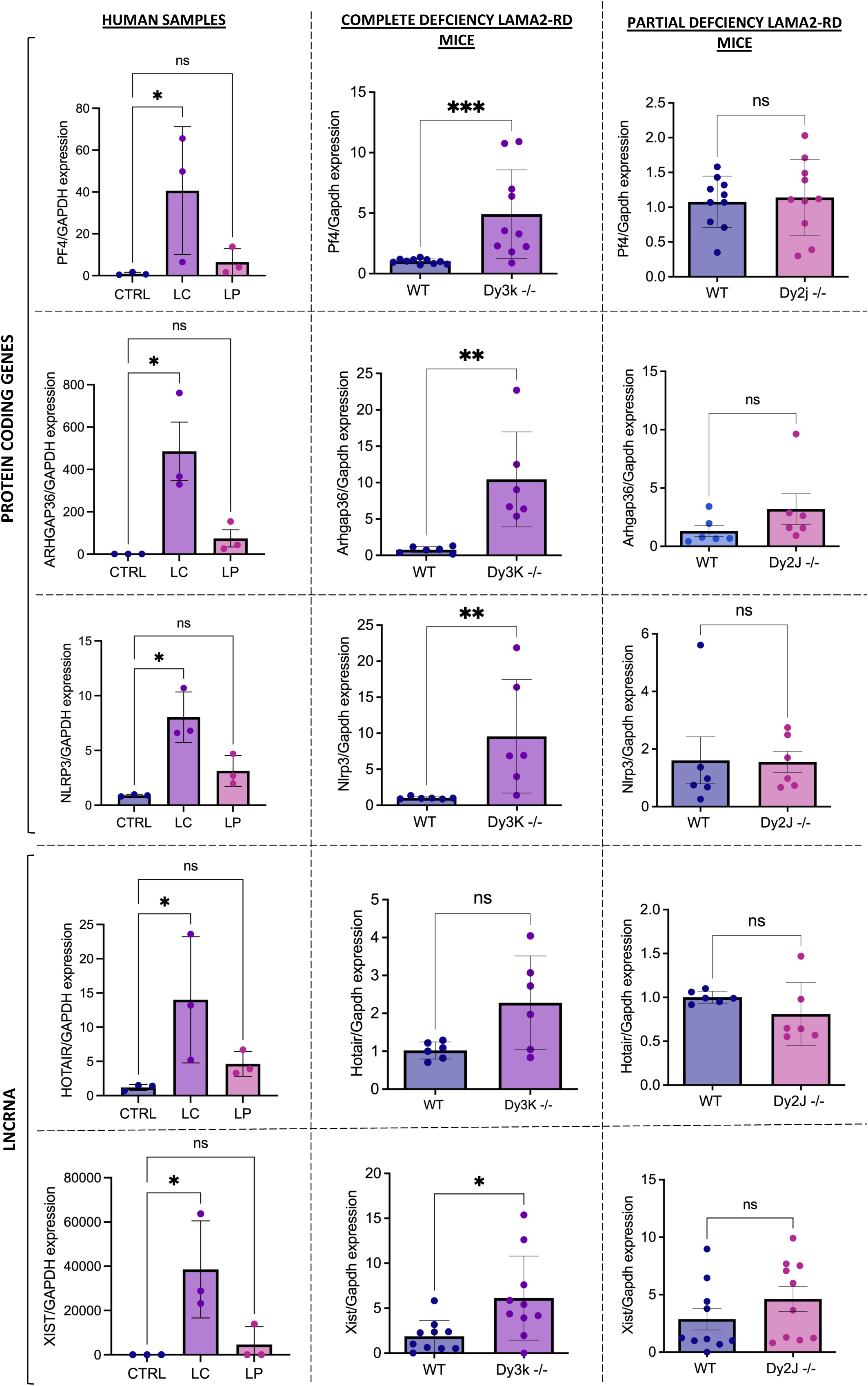
Novel LAMA2-RD biomarkers similarly altered in patients and mice models of Complete and Partial protein deficiency. Expression of selected targets validated both in sequenced human biopsies (n=3 samples/cohort) and muscle samples from LAMA2-RD mouse models (n=6-10 samples/cohort). Samples from the severe Dy^3k^/Dy^3K^ strain were collected at postnatal day (p) 28, corresponding to advanced disease stage, while samples from the milder Dy^2J^/Dy^2J^ mice were collected from adult animals (p60) in which the disease was fully established. q-values of pairwise comparisons in human diseased samples compared to healthy controls were all below 0.05 (*). P-values obtained from comparisons of wild-type (WT) and affected animals were indicated as (*) for p≤0.05, (**) for p≤0.01, (***) for p ≤ 0.001 and (ns) for p> 0.05. Data are expressed as means ± SD.

Finally, to verify whether the expression of selected genes was conserved across species, we examined by qPCR their expression in samples from vastus lateralis muscles of Dy^3K^/Dy^3K^ and Dy^2J^/Dy^2J^ dystrophic mice (animal model of Complete and Partial laminin-211 deficiency, respectively). While *Ostn, Col22a1* and *Comp* were not altered (or followed a different expression trend) in dystrophic animals (*Supplementary Figure 3a*), *Nlrp3, Arghap36, Pf4*, *Hotair* and *Xist* followed the same expression trend seen in human transcriptome data (*Figure 6, middle and right panel*), suggesting their potential as universal biomarkers of disease severity. Interestingly, *Xist* expression was not only altered in in Dy^3K^/Dy^3K^ females, but also in male mice (*Supplementary Figure 3b*), possibly hinting at a role beyond X-chromosome inactivation in dystrophic pathology.

## Discussion

LAMA2-RD is a rare neuromuscular disorder caused by absent or defective laminin-211, with clinical severity typically correlating with residual protein levels in skeletal muscle. While previous studies reported some molecular alterations in mild and severe phenotypes (Millino *et al.,* 2006), a comprehensive description of the muscle transcriptomic landscape defining each patient cohort is still missing. In our study, we relied on complementary bioinformatic tools to perform an integrated analysis of transcriptome data from patients with either Complete or Partial laminin-211 deficiency, identifying both shared and divergent molecular alterations and exploring their relevance to disease pathogenesis. By integrating gene enrichment, canonical pathway analysis, and upstream regulator prediction with focused gene-by-gene literature curation, we detangle the contribution of specific molecular mechanisms to disease pathogenesis in relation to laminin-α2 expression levels. Processes like fibrosis, chronic inflammation, disrupted homeostasis, altered cell cycle, metabolic dysfunction and oxidative stress were consistently altered in both cohorts, but were more severely perturbed in Complete cohort (reflecting the broader impact of full laminin-211 loss). Most of these pathways are commonly involved across muscular dystrophies affecting ECM components, membrane or cytoplasmic protein that disrupt homeostasis along the ECM-sarcomere axis (de las Heras *et al*., 2023; González-Jamett *et al*., 2022; Ignatieva *et al*., 2020; Segatto *et al*., 2020; Zanotti *et al*., 2023).

However, fibrosis has gained considerable attention as the key molecular driver of LAMA2-RD pathology rather than being a secondary consequence of tissue damage. Previous studies confirmed indeed that fibrosis, mediated by the joint action of TGF-β and renin-angiotensin signalling, is already prominently established in early postnatal muscle (both in patients and animal models), potentially arising from laminin-211 absence both in myocytes and tissue-resident fibroblasts (Accorsi *et al*., 2020). Except for a few genes (including *POSTN, SPP1* and *GPC3* which were much more expressed in Complete group), the majority of genes coding for ECM components (such as *COL1A1*, *COL1A2, ELN, FN1*) were similarly overexpressed in both cohorts, reinforcing matrix remodelling as a core pathological feature irrespective of residual laminin-211.

Nonetheless, growing evidence highlighted the importance of ECM-mitochondria crosstalk, demonstrating that ECM changes contribute to mitochondrial dysfunction and initiate stress responses acting on TGF-β release and SMAD signalling activation, thus creating a circle that interconnects fibrosis and metabolism (Piñeiro-Llanes *et al*., 2025; H. Zhang *et al*., 2024). Supporting this, we found that *PPARGC1A* metabolic hub was the most downregulated master regulator in both Cohorts. Importantly, most genes related to metabolic dysfunction including mitochondrial biogenesis, β-oxidation and electron transport chain were almost exclusively differentially expressed in the Complete cohort, similarly to what we found in many genes affecting carbohydrates and amino-acids metabolism. These findings indicate a broader metabolic deficit in laminin-211 deficient muscle which is more severely affected in the Complete absence cohort, in turn exacerbating cellular stress and impairing homeostasis. The broader deregulation of oxidative stress genes, including those involved in stress-dependent mechanisms like autophagy, ubiquitin-proteasome complex activation, and eventually cell death provides further evidence on the dysregulated pathway of this condition.

It’s worth highlighting that metabolic dysfunction further contributes to inflammation, specifically through the activation of inflammasomes in skeletal muscle, innate immune system cells and fibroblasts (Postma, 2020)(Elliott *et al*., 2018)(Irazoki *et al*., 2023; Y. Liu *et al*., 2020). In this regard, inflammasome-related genes including *NLRP3* were only increased in Complete cohort, outlining that this process is uniquely overactive in absence of laminin-211. *NLRP3* upregulation was also previously reported in 7 days old Dy^W^/Dy^W^ pups (a severely affected LAMA2-RD mouse model with minimal laminin-211 traces, considered equivalent to complete absence model) (Jeudy *et al*., 2011), providing further evidence of the early occurrence of inflammation in the pathogenesis of LAMA2-RD. Pyroptosis (a caspase-mediated form of inflammatory cell death process tightly linked to inflammasome which proved to be a valuable target in *mdx* mice (Dubuisson *et al*., 2022) was also exclusively increased in the Complete cohort, as were genes supporting the production of immune cells and the recruitment of inflammatory cytokines to damaged tissues. Such pronounced inflammatory microenvironment might be tied to the immunomodulatory role of laminin-211 itself. In fact, while our data clearly point to a role of laminin-211 deficiency in muscle and tissue-resident fibroblasts, *LAMA2* is also expressed in thymus; laminin-211 absence in this organ development and postnatal life was reported to result in selective thymocytes death (Iwao *et al*., 2000) which, in turn, was shown to remotely impair satellite cells proliferation and facilitate muscle aging (Zheng *et al*., 2022). Also, reduced *LAMA2* expression was linked to activated CD4 memory T cells in inflammatory milieu (Y. Jin *et al*., 2023), possibly increasing CD4+ T-cell infiltration and secretion of immunomodulatory cytokines that aggravate local inflammation.

Given laminin-211 crucial role in providing structural support and signal transduction through α-dystroglycan and α7β1 integrin binding, we expected these two functions to be also impacted by laminin-211 defects/absence. While both receptors contribute to skeletal muscle homeostasis through sarcoplasmic organization and mechanical force transmission (Boppart *et al*., 2006; Han *et al*., 2009), α-dystroglycan predominantly fulfils structural roles via the binding of dystrophin and, indirectly, to cytoskeletal actin, whereas integrin α7β1 acts primarily as a signalling hub for FAK, ILK and MAPK/ERK pathways, among others (Khyrul *et al*., 2004; J. Liu *et al*., 2008). The downregulation (although not dramatic) of genes involved in sarcoplasm remodelling and sarcomere function (including *NEB, TNNI1, TNNT1, TNNC1, TPM3, TTN*) likely reflects the lost/impaired laminin connection also with α-dystroglycan. Conversely, the dysregulation of cyclins, cyclin-dependent kinases (CDKs) and checkpoints (particularly in G1-S-G2-M transitions) might instead originate from the disrupted/missing link with integrins (Assoian & Schwartz, 2001; Kamranvar *et al*., 2022). Integrins also indirectly affect genes as *RB1* (predicted to be one of the downregulated upstream regulators in Complete cohort)(X. Zhu *et al*., 1996), thus affecting cell cycle progression. Genes linked to G2-M transition were downregulated in Complete but slightly upregulated in Partial cohort, likely due to residual laminin function. The altered expression of genes involved in mitotic spindle formation and chromosome segregation, often dependent on microtubules and cytoskeletal actin filaments, further supports impaired cell cycle progression. Furthermore, laminin deficiency results in a hostile niche for satellite cells impairing their ability to complete cell cycle and their regenerative capacity, indirectly contributing to muscle degeneration and fibrosis. Altered genes in Wnt, Notch and Hedgehog pathways suggested ongoing regeneration attempts in both cohorts (Bi *et al*., 2016; Norris *et al*., 2023; K. Zhang *et al*., 2018). Interestingly, while aiding muscle regeneration (Piccioni *et al*., 2014), Sonic hedgehog (Shh) signaling was also lately implicated in fibrosis and myofibroblast activation in multiple tissues (Gu *et al*., 2023)(H. Cao *et al*., 2020)(Fan *et al*., 2023) (Horn *et al*., 2012). Indeed Zhu *et al*. showed that *LAMA2* knockdown in mesenchymal cells elevates Shh expression, altering their commitment (Y. Zhu *et al*., 2020). Also *ARHGAP36* (among the most upregulated genes in both cohorts) activates Shh signaling and genes related to ECM production (Melo *et al*., 2023), supporting its role in disease pathology.

Our gene-tailored analysis shed light also on *PF4,* whose prominent upregulation restricted to Complete cohort interconnects most molecular events dysregulated in this patient group; PF4 has indeed gained attention as a key target in several fibrotic contexts taking part not only to myofibroblasts transition through TGFβ smad2/3 signaling and *ANGII* expression, but also to pyroptosis (Capitanio *et al*., 2024)(Wei, Wang, *et al*., 2024)(Wei, Peng, *et al*., 2024) licensing *NLRP3* transcription and IL-1β secretion by monocytes, thus contributing to inflammasome activation (Rolfes *et al*., 2020). Despite balanced sex ratios across diseased cohorts, also the lncRNA *XIST* was among the top upregulated genes in Complete but not Partial patients. Beyond X-chromosome inactivation (Markaki *et al*., 2021), *XIST* was recently described as a major regulator of ECM homeostasis and fibrosis via the miR-29b-3p/*COL1A1* axis (Yang et al., 2019)(W. Cao & Feng, 2019). Recent findings in osteoarthritis patients also linked low *LAMA2* expression with increased *XIST* (Huang *et al*., 2023). Importantly, in fibrotic contexts, *XIST* upregulation was also shown to affect cell cycle by increasing *CDKN1A* expression (as also seen in LAMA2-RD patients and animal models) (Yang *et al*., 2019) and to exacerbate both inflammation and apoptosis by acting on *NLRP3* (Ma *et al*., 2019), Toll-like receptor 4 and angiotensin II (Jin *et al*., 2019). Nonetheless, *XIST* overexpression was also associated to mitochondrial dysfunction and concomitant increase of reactive oxygen species (ROS) production in hepatocytes(Wu *et al*., 2023), suggesting that its dysregulation in dystrophic muscle could be also related to the more severe metabolic impairment observed in Complete cohort. Another lncRNA (*HOTAIR*) exclusively upregulated in Complete cohort promotes inflammasome activation and ROS generation, both by inhibiting the expression of the antioxidant protein NRF2(You *et al*., 2023) and by sponging miR-129-5p (Y. Wang *et al*., 2022). As for *XIST*, *HOTAIR* targets *NLRP3* amplifying the inflammatory response (Y. F. Liu *et al*., 2021) and leading to pyroptosis. Interestingly, *HOTAIR* expression is inversely correlated with the expression of *BDNF* (Deb *et al*., 2024), a neurotrophic factor exerting protective roles in dystrophic muscle (Kristensen *et al*., 2022), suggesting its potential in maintaining tissue homeostasis.

Given their involvement in many molecular processes altered in LAMA2-RD and in the respective murine models (and other muscular dystrophies), the above-mentioned genes/pathways may serve as universal biomarkers for disease stratification. While this study is limited by the small sample size due to the scarce availability of human biopsies, bulk transcriptome data offer a unique opportunity to study the intertwined molecular events shaping LAMA2-RD muscle pathology. By delineating unique marks tied to tissue-specific laminin-211 roles, our study presents a detailed landscape of the key biological features defining each patient cohort and opens new avenues for therapeutic interventions targeted to secondary pathological cascades in LAMA2-RD. These findings pave the way for future single-cell transcriptomics studies aimed at unravelling the contributions of distinct muscle-resident cells to disease progression.

## Methods

### Ethical Statement

The use of human biological samples was ethically approved and consent for research was obtained to facilitate pharmacological, gene and cell therapy trials in neuromuscular disorders (NRES Committee London-West London & GTAC, REC reference number 06/Q0406/33) and to allow the use of muscle biopsy to study pathogenesis and the expression and regulation of genes associated with neuromuscular disorders (NRES Committee London—REC reference 13/LO/1894), in compliance with national guidelines regarding the use of human-derived biological samples for research. Written informed consent was obtained from patient’s guardians.

### Patients’ muscle biopsy collection

Human samples selected for this study included quadriceps muscle biopsies from 4 LAMA2-RD patients with complete laminin-211 deficiency (severe disease), 4 patients with partial laminin-211 deficiency (generally milder severity) and 4 age-matched (0-3 years old) non-affected controls (individuals in whom a neuromuscular disease was excluded by the relevant investigations and/or samples obtained opportunistically during planned orthopedic surgery). Details regarding selected samples are indicated in the table below. LAMA2-RD biopsies were all obtained as part of standard clinical care for diagnostic purposes, and stored with consent at the MRC Centre for Neuromuscular Diseases Biobank (London, UK)(Ala *et al*., 2025). One LAMA2-RD muscle biopsy was shipped to the UK from San Raffaele Hospital Institute of Experimental Neurology (Milan, Italy), where it was obtained for diagnostic purposes.

| Sample ID | Sex | Age at biopsy | Laminin-211 | LAMA2 mutation(s) | Phenotype |
| --- | --- | --- | --- | --- | --- |
| CTRL_01 | M | 2 yr 1 mth | Normal | None | Healthy |
| CTRL_02 | M | 2 yr | Normal | None | Healthy |
| CTRL_03 | M | 5 yr | Normal | None | Healthy |
| CTRL_04 | F | 2 yr 8 mth | Normal | None | Healthy |
| LAMA2_COMPLETE_01 | F | 10 mth | Absent | Hom. c.736C>T(p.Gln246X) | Severe - never walked |
| LAMA2_COMPLETE_02 | M | 11 mth | Absent | Hom. c.1306+2T>G | Severe - never walked |
| LAMA2_COMPLETE_03 | F | 3 mth | Absent | Hom. c.4690C>T (p.His1564Tyr) | Severe - never walked |
| LAMA2_COMPLETE_04 | F | 2 yr | Absent | Hom. c.2049_2050del (p.Arg683Serfs*21) | Severe - never walked |
| LAMA2_PARTIAL_01 | M | 4 yr | Reduced | Het. c.2749+2_3insTT (intronic) and<br>Het. c.6443delT (p.Val2148fs) | Mild - ambulant |
| LAMA2_PARTIAL_02 | F | 1 yr 5 mth | Traces | Het. c.1303C>T (p.Arg435X) and<br>Het. c.155T>A (p.Ile52Asn) | Severe - never walked |
| LAMA2_PARTIAL_03 | F | 1 yr 4 mth | Reduced | Het. c.5562+5G>C (intronic),<br>Het. c.9211+6C>T (intronic, same allele)<br>and Het. c.3976C>T (p.Arg1326X) | Intermediate -<br>ambulation lost over<br>time |
| LAMA2_PARTIAL_04 | F | 4 yr | Reduced | Not disclosed | Mild - ambulant (but<br>delayed motor<br>milestones) |

### RNA extraction from human skeletal muscle tissue

Total RNA was extracted from selected snap frozen muscle biopsies by using the RNeasy Mini kit (QIAgen, Ltd., Manchester, UK) according to manufacturer’s instructions. RNA quality was assessed by the UCL Bioinformatics Core office via the Agilent 2200 TapeStation instrument to ensure that all samples submitted for sequencing had an RNA Integrity Number (R.I.N) equal to or above 6.

### RNA sequencing and raw data processing

150 ng total RNA was shipped to the Center for Mendelian Genomics (Broad Institute of MIT and Harvard), where sequencing library was prepared using the Truseq Strand Specific Large Insert kit and sequenced with Illumina instrument, with a depth of coverage of 75 million 101 bp paired-end reads. Sequencing reads were aligned to the reference genome (hg38 Ensembl) by running STAR v2.6.1d(Dobin *et al*., 2013). Principal components analysis (PCA) was performed on normalized (transcripts-per-million) and log-transformed expression values, with GTEx v8 muscle and adipose tissue samples included for comparison. Differential expression analysis was done in the R programming language using the DESeq2 package (Love *et al*., 2014) for reads data normalization and statistical analysis. DESeq2 outputs of differentially expressed genes (DEGs) include the fold changes of expression indicated as base2 logarithm (or Log2FoldChange) and scores indicating the statistical significance, i.e. p-values and p-values adjusted for multiple comparisons (q-values). Genes with a q-value equal to (or minor than) 0.05 were considered for further analysis, in accordance with DESeq2 guidelines. Given the limited number of available muscle samples, each analysis presented in our work was conducted considering all significant dysregulated genes, including those with an expression fold change between -2 and +2.

### Gene Enrichment analysis

ShinyGO v0.80 webtool was used to obtain functional annotations of DEGs and predict enriched Gene Ontology (GO) terms related to cellular components, molecular function and biological processes (Ge *et al*., 2020). Altered biological processes were further investigated by querying the most significant Molecular Signatures Database (MSigDB) hallmarks (Liberzon *et al*., 2015). False Discovery Rate (FDR) cut-off was set at 0.5, while over-represented entries were represented by a fold enrichment bigger than (or equal to) 1.

### Bioinformatic analysis of dysregulated pathways and upstream regulators

The thorough investigation of perturbed molecular mechanisms was performed by relying on the Ingenuity Pathway Analysis software (IPA) from Qiagen. IPA uses its knowledge-based intelligence to run a Core analysis in which DEGs are grouped in its collection of curated canonical pathways, generating molecular networks and inferring upstream regulators of the submitted gene dataset (Krämer *et al*., 2014). The probability of pathway activation or inhibition was expressed by a score (z-score) which also reflects the number of DEGs matching the pathway. P-value was calculated by the right-tailed Fisher’s Exact test while the statistical significance for canonical pathway analysis, indicated as log(p-value), was set at values equal to (or major than) 1.3. When considering predicted upstream regulators, the p-value of overlap cut-off was set at 0.05.

### Mice maintenance and muscle tissue isolation

All the experiments performed on dystrophic LAMA2-RD mice received ethical approval and were performed in agreement with the Ospedale San Raffaele Institutional Animal Care and Use Committee (IACUC authorization #1086). The Dy^2J^/Dy^2J^ mice were purchased from Jackson Laboratories (Bar Harbor, USA), while Dy^3K^/Dy^3K^ mice were previously described in (Miyagoe *et al*., 1997). Both strains were maintained in the C57BL/6J background. Genotyping was performed by isolating genomic DNA from tail biopsies using DirectPCR solution (Viagen), according to manufacturer’s instructions. Primer sequences used for genotyping each strain are indicated below.

Dy^2J^/Dy^2J^ forward primer:5’-CTCTATTACTGAACTTTGGATG-3’ Dy^2J^/Dy^2J^ reverse primer: 5’-TCCTGCTGCCTGAATCTTG-3’ Dy^3K^/Dy^3K^ forward primer: 5’-CAGGTGTTCCAGATTGCC-3’ Dy^3K^/Dy^3K^ Neo primer: 5’-CCCGTGATATTGCTGAAG-3’

Dy^3K^/Dy^3K^ reverse primer: 5’-CCTCTCCATTTTCTAAAG-3’

Mice were euthanized by CO2 inhalation. Quadriceps muscle isolated from sacrificed animals was snap frozen in liquid nitrogen and stored until needed for RNA isolation.

### RNA extraction from animal skeletal muscle tissue

Total RNA was extracted from quadriceps muscle using TriPure Isolation Reagent (Roche) according to the manufacturer’s instructions. Briefly, muscles were homogenized in the presence of TriPure Isolation Reagent, total RNA was isolated from the homogenate with chloroform and then precipitated with isopropanol. The RNA pellet was finally resuspended in RNAse-free water.

### Retrotranscription and qRT-PCR

The same protocol was used to retrotranscribe human and animal RNA. One microgram of total RNA was reverse-transcribed using the High-Capacity cDNA Reverse Transcription Kit (Applied Biosystems), following the manufacturer’s instructions. Quantitative reverse transcriptase PCR was run on the Applied Biosystems 7900HT Real-Time PCR System. The reaction was set up using the 2× TaqMan PCR Master Mix (Applied Biosystems) as recommended by the manufacturer. qRT-PCR was run following the thermocycling program: 95 °C for 3 minutes, and 40 cycles of 10 seconds at 95 °C and 1 minute at 60 °C. The PrimeTime qPCR Std® and TaqMan® assays used to confirm gene expression levels are described in tables below. The expression of each gene of interest was normalized to that of *GAPDH.* Data analysis was performed with the comparative cycle threshold (ΔΔCt) method (Livak & Schmittgen, 2001).

| qRT-PCR assays - HUMAN |  |  |  |
| --- | --- | --- | --- |
| Gene_ID | qPCR Assay name | Manufacturer | Dye |
| *ARHGAP36* | Hs00376478_m1 | Thermo Fisher Scientific | FAM |
| *COL22A1* | Hs01377218_m1 | Thermo Fisher Scientific | FAM |
| *COMP* | Hs00164359_m1 | Thermo Fisher Scientific | FAM |
| *GAPDH* | Hs02786624_g1 | Thermo Fisher Scientific | FAM |
| *HOTAIR* | Hs05502358_s1 | Thermo Fisher Scientific | FAM |
| *NLRP3* | Hs00918082_m1 | Thermo Fisher Scientific | FAM |
| *OSTN* | Hs00898258_m1 | Thermo Fisher Scientific | FAM |
| *PF4* | Hs00427220_g1 | Thermo Fisher Scientific | FAM |
| *XIST* | Hs01079824_m1 | Thermo Fisher Scientific | FAM |

| qRT-PCR assays - MICE |  |  |  |
| --- | --- | --- | --- |
| Gene_ID | qPCR Assay name | Manufacturer | Dye |
| *Arhgap36* | Mm.PT.58.13567184 | Integrated DNA Technologies | FAM |
| *Col22a1* | Mm.PT.58.12878924 | Integrated DNA Technologies | FAM |
| *Comp* | Mm.PT.58.21638867 | Integrated DNA Technologies | FAM |
| *Gapdh* | Mm.PT.39a.1 | Integrated DNA Technologies | FAM |
| *Hotair* | Mm.PT.58.30586279 | Integrated DNA Technologies | FAM |
| *Nlrp3* | Mm.PT.58.13974318 | Integrated DNA Technologies | FAM |
| *Ostn* | Mm.PT.58.11223807 | Integrated DNA Technologies | FAM |
| *Pf4* | Mm.PT.58.32040678.g | Integrated DNA Technologies | FAM |
| *Xist* | Mm.PT.58.8416342 | Integrated DNA Technologies | FAM |

### Statistical Analysis

Statistics used by each bioinformatic software to interpret RNA-sequencing data are indicated in the corresponding paragraph. Gene expression data obtained by qRT-PCR were analyzed with GraphPad Prism v8.0 software by applying nonparametric statistical tests, given small sample size and non-Gaussian distribution. Specifically, Mann-Whitney and Kruskal-Wallis tests were applied to infer significance when comparing two and three groups, respectively, with p-value cut-off set at 0.05. For multiple pairwise comparisons (i.e., complete and partial groups versus control), p-values were adjusted (q-value) using Dunn’s correction. A q-value < 0.05 was considered statistically significant.

## Acknowledgements

We kindly acknowledge funding received from CureCMD (grants awarded in 2018, 2020 and 2022), and the partial support of funds received in 2024 both from LifeArc/MDUK (as part of the Muscular Dystrophy Translational Research Fund) and Marie Curie Postdoctoral Fellowship (Project 101149955). The figures from section “the paper explained” were generated using BioRender, while pathway figures were generated with Ingenuity Pathway Analysis software (licensed by the UCL Genomics facility). The authors are also grateful for the support received from the Neuromuscular Biobank and the National Institute for Health and Care Research Biomedical Research Centre at Great Ormond Street Hospital for Children, NHS Foundation Trust, University College London, and the Neuromuscular Repair Unit of Ospedale San Raffaele. We also acknowledge support received from Prof. Daniel MacArthur, Prof. Anne O’Donnell-Luria and the whole team of the Center for Mendelian Genomics at the Broad Institute of Harvard and MIT. Finally, we wish to thank Dr. Rita Barresi for providing human tissue included in the study, and Dr. Sara Aguti and Dr. Elena Marrosu for their help in processing human samples.

## Author contributions

Study conceptualization: Veronica Pini, Francesco Muntoni; RNA-sequencing data processing: Ben Weisburd; Bioinformatic analyses: Ben Weisburd, Anne, Veronica Pini; Processing of human muscle samples and qPCR: Francesco Catapano; Collection of murine samples and qPCR: Rosa Bonaccorso; Statistical analysis: Veronica Pini, Francesco Catapano, Rosa Bonaccorso; Data analysis and interpretation: Veronica Pini, Francesco Muntoni, Stefano C. Previtali; Manuscript writing-original draft: Veronica Pini; Manuscript review and editing: Francesco Muntoni, Stefano C. Previtali, Francesco Catapano; Funding acquisition: Veronica Pini, Francesco Muntoni.

## Data availability

The datasets produced in this study will be made available in the Gene Expression Omnibus (GEO) repository pending approval of the manuscript.

## Supplementary figures legend

**Supplementary Figure 1:**
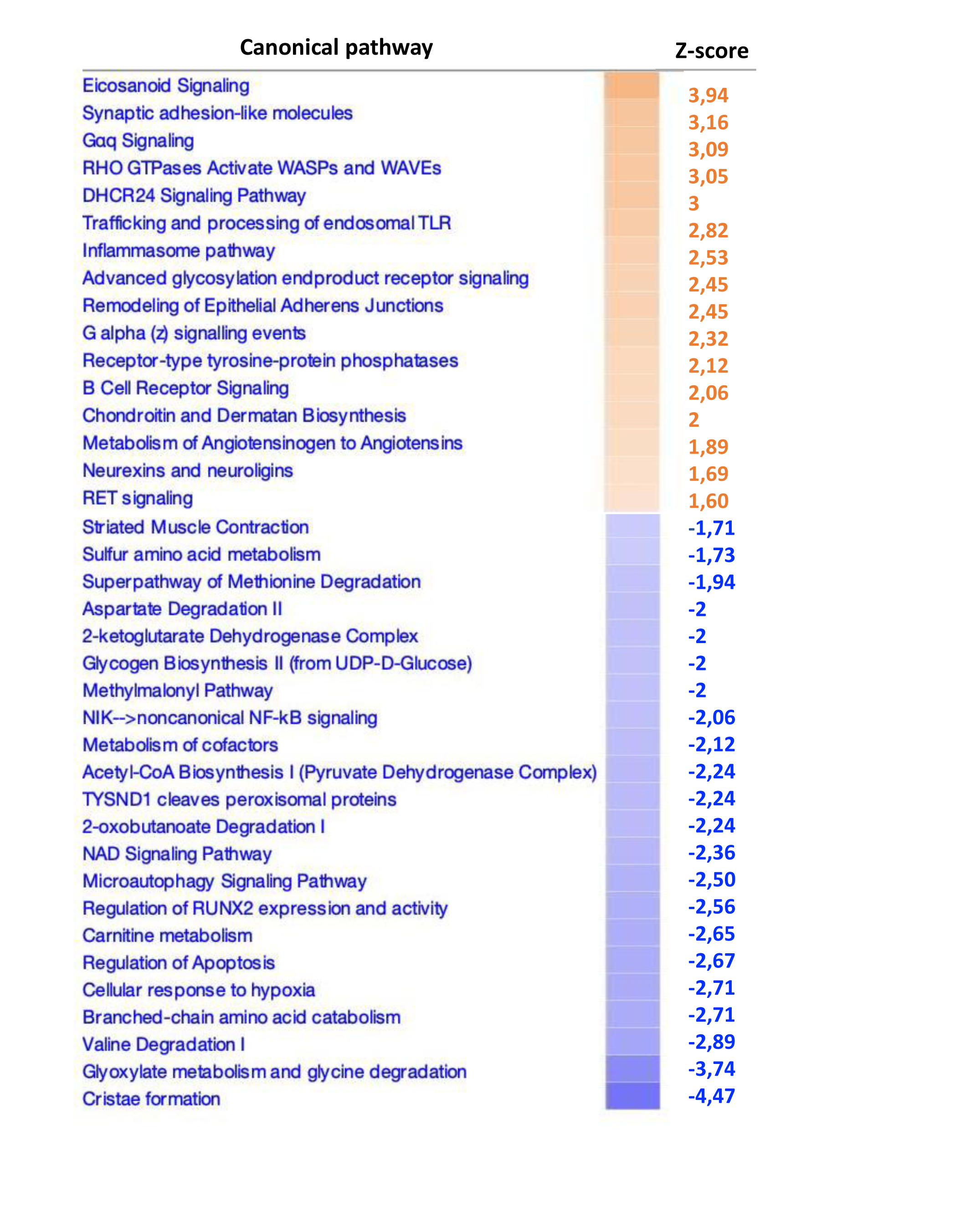
canonical pathways only dysregulated in the Complete cohort (z=+-1.5) Canonical pathways are ordered according to descending z-score values.

**Supplementary Figure 2:**
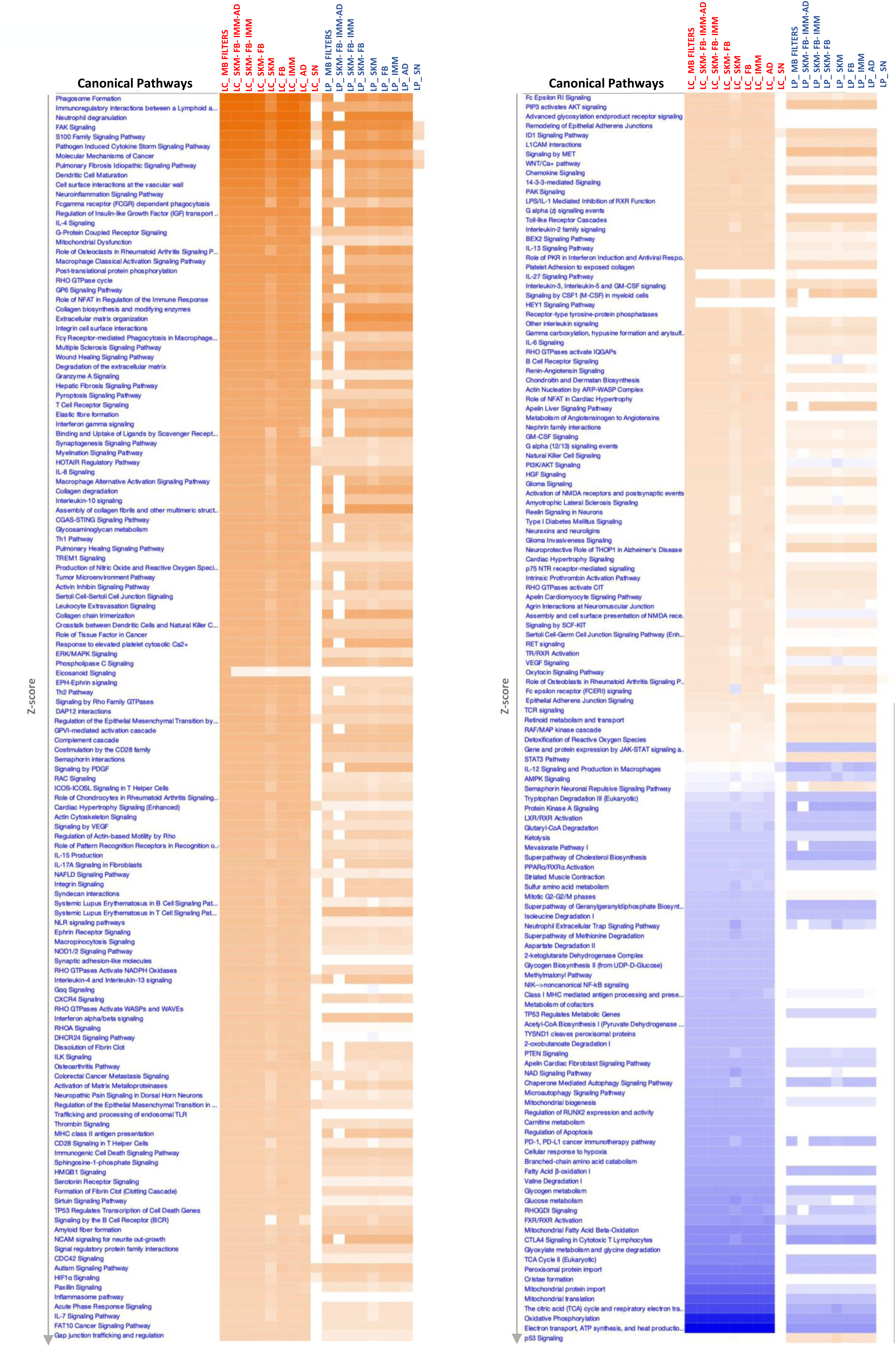
Comparison of canonical pathways altered in LAMA2-RD patients with Complete (LC) and Partial (LP) laminin-211 deficiency using Ingenuity Pathway Analysis (IPA) Canonical pathways identified as significantly altered in each LAMA2-RD patient cohort, filtered by z-score. For each patient cohort, the first core analysis (MB filters) includes findings observed in skeletal muscle and potential contaminants such as endothelial cells, chondrocytes, fibroblasts, immune cells, adipocytes and sciatic nerve. Serial core analyses were run based on increased filter stringency (i.e. limiting findings against one or more of the above-mentioned cell types). SKM = skeletal muscle; FB= Fibroblasts; IMM = Immune cells; AD = adipocytes; SN= sciatic nerve.

**Supplementary Figure 3:**
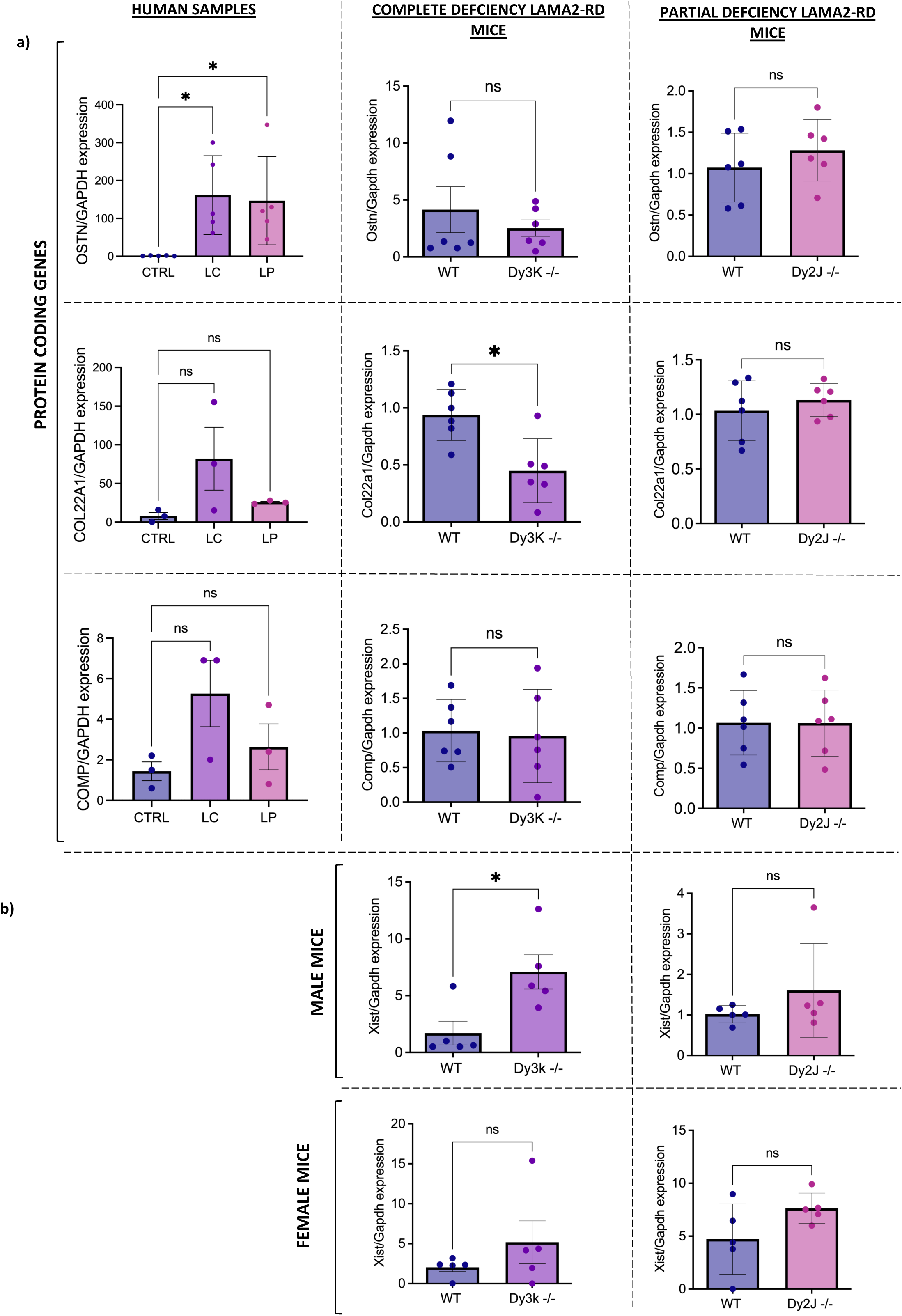
Divergence between murine models and human qPCR validation. a) Genes selected from human sequenced samples (n=3 samples/cohort) with discrepant expression in muscle tissues isolated from Dy^3K^/Dy^3K^ (p28) and Dy^2J^/Dy^2J^ (p60) mice (n=5-6/cohort); 1b) *Xist* expression separately assessed in female and male LAMA2-RD murine models. *=p≤0.05; ns=not significant (p> 0.05). Data are expressed as means ± SD.

**Supplementary Table 1:** DESeq2 dataset showing all differentially expressed genes in each LAMA2-RD cohort compared to healthy controls.

**Supplementary Table 2:** Altered canonical pathways in each cohort returned by Ingenuity Pathway Analysis

**Supplementary Table 3:** z-scores obtained upon performing multiple comparison analysis in Ingenuity

**Supplementary Table 4:** Targeted gene analysis grouped according to their main biological role

